# Selection following gene duplication shapes recent genome evolution in the pea aphid *Acyrthosiphon pisum*

**DOI:** 10.1101/643544

**Authors:** Rosa Fernández, Marina Marcet-Houben, Fabrice Legeai, Gautier Richard, Stéphanie Robin, Valentin Wucher, Cinta Pegueroles, Toni Gabaldón, Denis Tagu

## Abstract

Ecology of insects is as wide as their diversity, which reflects their high capacity of adaptation in most of the environments of our planet. Aphids, with over 4,000 species, have developed a series of adaptations including a high phenotypic plasticity and the ability to feed on the phloem-sap of plants, which is enriched in sugars derived from photosynthesis. Recent analyses of aphid genomes have indicated a high level of shared ancestral gene duplications that might represent a basis for genetic innovation and broad adaptations. In addition, there is a large number of recent, species-specific gene duplications whose role in adaptation remains poorly understood. Here, we tested whether duplicates specific to the pea aphid *Acyrthosiphon pisum* are related to genomic innovation by combining comparative genomics, transcriptomics, and chromatin accessibility analyses. Consistent with large levels of neofunctionalization, we found that most of the recent pairs of gene duplicates evolved asymmetrically, showing divergent patterns of positive selection and gene expression. Genes under selection involved a plethora of biological functions, suggesting that neofunctionalization and tissue specificity, among other evolutionary mechanisms, have orchestrated the evolution of recent paralogs in the pea aphid and may have facilitated host-symbiont cooperation. Our comprehensive phylogenomics analysis allowed to tackle the history of duplicated genes to pave the road towards understanding the role of gene duplication in ecological adaptation.

## Introduction

Aphids are insect pests belonging to the order Hemiptera, which diverged some 280-250 million years ago. They feed exclusively on plant phloem sap, a trait that involved specific adaptations such as an obligatory symbiosis with bacteria of the genus *Buchnera*, which supplies aphids with essential amino acids that are missing in the phloem sap. In addition, to adapt to stressful environments such as cold, predation and parasitism (Vellichirammal et al. 2016), aphids have developed several plastic phenotypic traits, involving winged and apterous morphs, or sexual oviparous and parthenogenetic viviparous morphs. Although several studies have addressed the genetic mechanisms of these adaptations at the molecular level, the evolutionary forces underlying these genomic changes are still poorly understood. Today, several aphid genomes are publically available and all show a high level of gene duplication and expansions (The International Aphid Genomics Consortium 2010; Mathers et al. 2017;Li et al. 2019). Some of these duplications are shared between aphid species, but most of them are lineage-specific (IAGC 2010). The modes of evolution of gene duplicates occurring in these species are not yet fully determined. Whether or not duplicated or expanded gene families are in relation with the above-mentioned or other functional innovations enabling adaptive evolution in aphids is still largely unknown (Huerta-Cepas et al. 2010; Simon et al. 2011).

There are at least four different outcomes for gene duplicates (reviewed in Capella-Gutierrez et al. 2009; Innan and Kondrashov 2010). First, while one duplicate keeps the original function, the other acquires a new function (neofunctionalization). Second, each of the two duplicated genes keeps part of the functions of the ancestral gene, so that they jointly cover the original functions (subfunctionalization). Third, when the increase in gene dosage is beneficial, the two copies are maintained in the absence of functional divergence. And fourth, the most common output of gene duplication is the inactivation by accumulation of mutations of one of the duplicated genes (pseudogenization). Several evolutionary forces can drive these different outcomes, for instance relaxed selection for subfunctionalization, purifying selection for neofunctionalization or deleterious mutations for pseudogenization (Lynch and Conery 2000; Han et al. 2009; Innan and Kondrashov 2010). These different scenarios can be addressed by scrutinizing patterns of variation of gene families including lineage-specific duplications (Han et al. 2009; Innan and Kondrashov 2010; Pegueroles et al. 2013; Pich I Roselló and Kondrashov 2014).

More recently, sub- or neofunctionalization have started to be assessed by epigenetic regulation (Robin and Riggs 2003). Acquiring and losing functions can occur, among other means, by modification of chromatin states, which drive the transcriptional activities of genes. The so-called ‘open chromatin’, in which accessible DNA allows for active transcription, can be opposed to the so-called ‘‘closed’ chromatin, which is compact and transcriptionally repressed. Little is known about the role of chromatin in determining the fate of duplicated genes, but it is intuitive to think that two duplicated gene copies could have spatially or temporally different chromatin states, thus resulting in different transcription patterns. For instance, Keller and Yi (2014) showed that the DNA methylation of gene promoters of both copies of young duplicates in humans is higher than that of old duplicates. This observation stands for different tested tissues, indicating that this trait is not related to tissue-specificity regulation, as DNA methylation is known to regulate transcription. Thus, it could be hypothesized that chromatin state influences the expression of duplicated copies - and thus consequently their evolution - possibly as a protection against possible misregulations by dosage compensation (Chang and Liao 2012), before mutations occur and genetic selection operates. It is worth noting that divergent epigenetic environments may result in subfunctionalization (e.g. through changed expression patterns), but the epigenetic differences themselves do not result from functional differences between the copies.

Here, we test the hypothesis that gene duplication - particularly recent duplicates - in the pea aphid *Acyrthosiphon pisum* is a source of innovation fueled by selection. For this, we anchor our study on a phylogenomic approach exploring for the first time ten hemipteran genomes, including six aphid species. We show that (i) a large proportion of gene duplications are under positive selection in *A. pisum* and affect a large number of biological functions (most notably oo- and morphogenesis and host-symbiont cooperation), (ii) asymmetrical rates of young paralogs coupled to positive selection suggest neofunctionalization is a main force reshaping the pea aphid genome, (iii) a third of young duplicates show divergent tissue expression patterns, consistent in some cases with subfunctionalization by tissue specialization and others with neofunctionalization through gain of gene expression, and (iv) chromatin accessibility of the transcription start site (TSS) can change between genes in duplicated gene pairs, although it cannot directly explain their transcriptional state in *A. pisum*.

## Results and Discussion

### 1. Young paralogs in *A. pisum* are under neofunctionalization and involve diverse biological functions

We built a phylome (*i.e.* the complete collection of phylogenetic trees for each gene encoded in a genome) for *A. pisum* in the context of hemipteran evolution, including five additional aphid species and three basal Sternorrhyncha species (Fig. 1A). The phylome was then scanned for the presence of species-specific duplications. A total of 5,300 species/specific duplication events were detected in the *A. pisum* phylome that were clustered in 1,834 paralogous families. Due to the complexity of analysing and interpreting highly expanded and old duplications, we divided the duplications into two different sets. The first set consisted of pairs of genes specifically duplicated in *A. pisum* which fulfilled the following criteria: i) they presented they presented a single-copy ortholog in at least two of the other species in the phylome, ii) the node where the two sequences duplicated in the tree was well supported, iii) the tree reconstructed only using the selected paralogs and orthologs had to retain the duplication event. A second dataset focused on genes that were duplicated more than once specifically in *A. pisum*. In the case of these larger gene family expansions only pairs of genes found at the tips were considered. These pairs also had to fulfill the previous requirements. A total of 843 duplication events containing 1686 genes were selected for further analysis. Among those, 606 came from single duplication events whereas the remaining 237 were extracted from larger duplication events. (see **Table S3** for the complete list of selected genes and families). Note also that in some cases (8.5% of cases) an ortholog to the closest relative to *A. pisum* (*Myzus persicae*) was missing and hence there is the possibility that these particular duplications are older.

**Figure 1.**
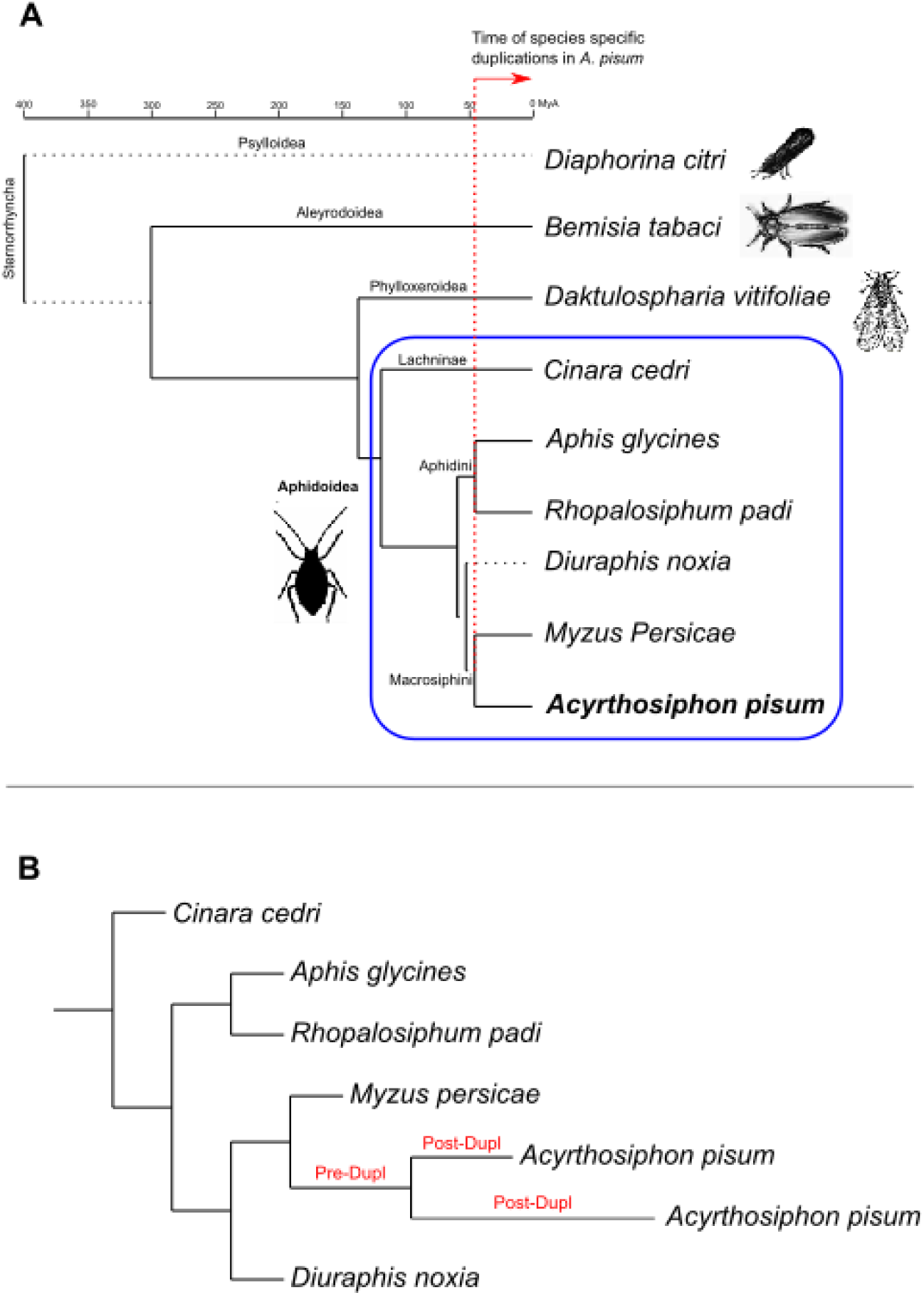
**A:** Chronogram of Sternorrhyncha interrelationships. Systematic classifications (superfamily, family, subfamily and tribe) are shown in each node/branch. Images selected from PhyloPic. Divergence times taken from TimeTree (Kumar et al. 2017). Dotted branches represent lineages for which divergence times were not available. **B:** Example of individual gene tree showing a duplication in *A. pisum*, as the genes selected from the present study (see Material and Methods). Pre- and post-duplication branches as defined for the positive selection analysis are highlighted in red.

We calculated the relative age of the selected duplications using the number of synonymous substitutions per synonymous site (dS) as a proxy. We estimated dS for each internal and terminal branch of each gene tree using the “free ratio branch model” from codeML and we filtered out specific genes with dS > 2 and dS < 0.01 (see Material and Methods for details). By comparing the distribution dS in each copy of the selected duplications (*A. pisum* Post-Dup) with the pre-duplication branches (*A. pisum* Pre-Dup) and single-copy orthologs (Fig. 1B), we showed that lineage-specific genes in *A. pisum* are enriched in recent duplications represented by their low dS values compared to other species (Fig. 2). Since it is known that gene conversion (GC) may decrease the divergence between paralogs, we scanned the respective coding sequences for the presence of GC tracts using GENECONV software (Sawyer 1989). We detected that 187 duplications (22.2%) showed evidence of a GC event between the two *A. pisum* sequences that remained significant after multiple-comparison correction. From those, 27 occurred in tandem duplicates (28.7% of total duplications in tandem), 31 in duplications in the same contig (31%) and 117 in duplications in different contigs (19.3%). Thus, tandem duplicates do not seem to be particularly enriched in GC, however we cannot rule out that some of the selected duplicates may be older than inferred due to gene conversion. In addition, due to the fragmentation of the genome assembly used it is possible that some tandem duplicates are not detected in our analyses. In an initial characterization of our set of recent gene duplications, we estimated the median identity for each protein sequence of each gene family alignment using trimAl v1.3. We observed that *A. pisum* duplicates were consistently (and significantly) less similar at the sequence level between them than when compared to single-copy orthologs, suggesting that their sequences are diverging faster (**Fig. S1A**). Thus, despite the presence of GC tracts, this process was not enough to homogenize the sequence of the gene duplicates. We discarded removing the GC tracts from the sequences because according to the literature DNA sequences are useful to detect the presence of GC but not to correctly infer their length (Mansai et al. 2011).

**Figure 2:**
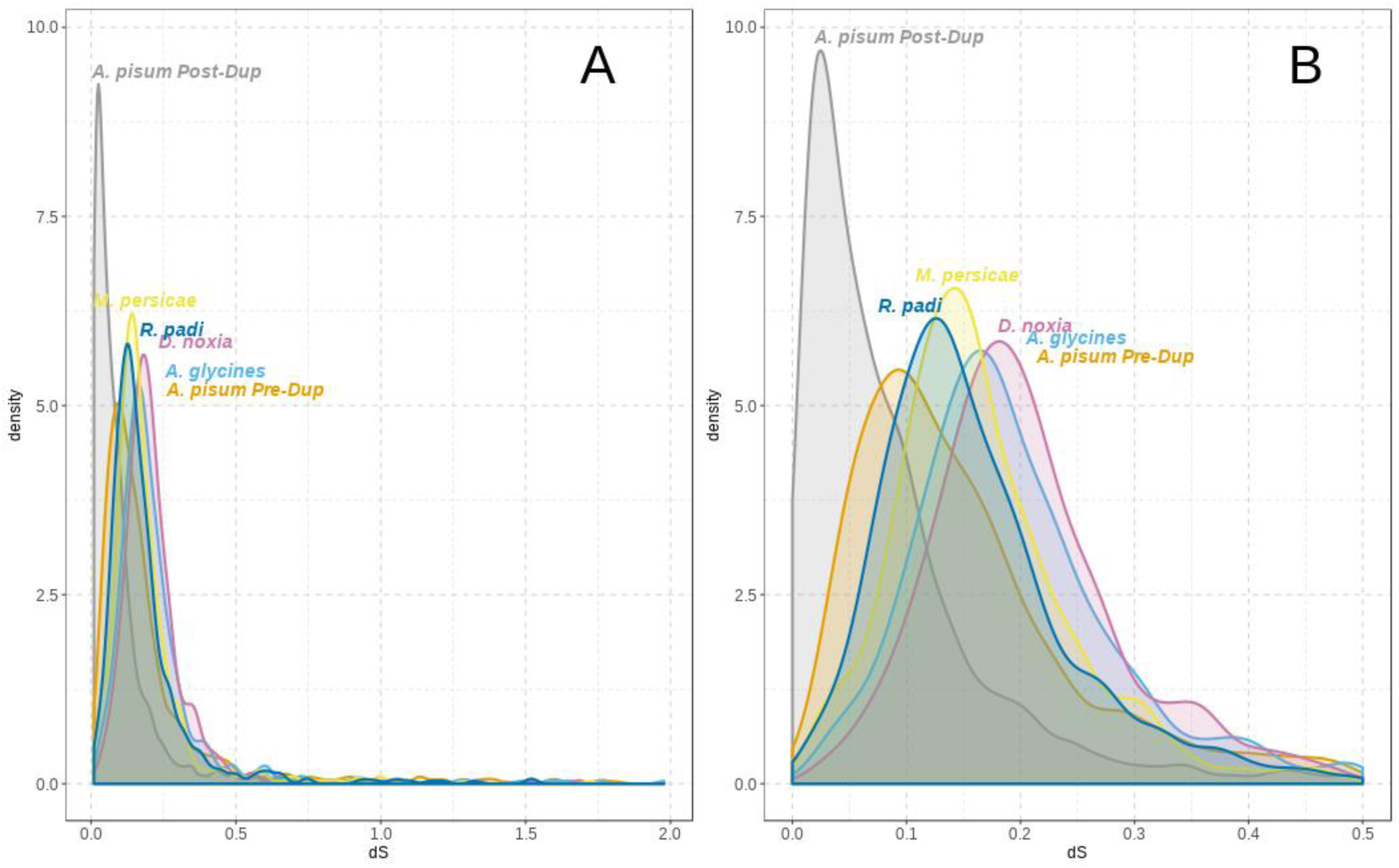
dS values for the selected duplications after filtering genes with dS >2 and dS <0.01 (**A**), and zoom by limiting x-axis to 0.5 (**B**). *C. cedri* was removed from this plot for visualization. dS for *A. pisum* was calculated before (*A. pisum* Pre-Dup) and after (*A. pisum* Post-Dup) the duplication took place. See Figure 1B for “Pre-Dup” and “Post-Dup” explanation.

To test the hypothesis of faster evolution of recently duplicated genes and to evaluate the pace of evolution in our set of recent gene duplications, we calculated the rate of evolution (dN/dS) for each gene family using codeML software from PAML package v4.9 (see Material and Methods for details). This software computes individual estimations for each branch of a given tree, allowing us to distinguish the evolutionary rate before and after the duplication (hereafter called as pre-duplication (Pre-Dup) and post-duplication (Post-Dup) branches, see Fig. 1B). To facilitate the interpretation of the results, gene duplicates were divided into two groups: “strict duplicates” *(i.e.*, genes with only two copies, ~72% of the selected duplicates) and “expansions” (genes with more than two copies, see Material and Methods for details). Paralogs of both “strict duplicates” and “expansions” had significantly faster rates as compared to their pre-duplicated ancestors as well as to single copy orthologs (Fig. 3A, **Fig. S2**). We then classified paralogous copies of each duplicated gene pair into “fast” and “slow” evolving copies, according to the dS values of the branch subtending each copy (see Material and Methods), which allows to distinguish between subfunctionalization and neofunctionalization scenarios (Sandve et al. 2018). Evolutionary rates were not homogeneous in the two copies, since the fast post-duplication copy is evolving more rapidly than both the slow post-duplication copy and the pre-duplication ancestor in the “strict duplicates” subset (Fig. 3B). The statistical significance of these differences in dN/dS values depends on the percentage of divergence between duplicate copies, reaching significance when pairs with lower than 50% divergence are discarded from the analysis, which correspond to ~1.4% of the duplicates that passed dS filters (Fig. 3C). The “expansions” subset showed the opposite pattern, in which the slow copy has higher dN/dS ratio than the fast (**Fig. S2B**). It is important noting that expansions are complex families, which are more prone to include missannotated genes or pseudogenes, which may influence our results. Thus, conclusions from this subset should be taken with caution. Asymmetrical evolution of gene duplicates has been observed in several organisms, such as fungi, *Drosophila melanogaster*, *Caenorhabditis elegans* and human, which was attributed to relaxed selective constraints and, in some cases, to the action of adaptive selection (Conant 2003; Zhang 2005; Scannell and Wolfe 2008; Pegueroles et al. 2013; Pich I Roselló and Kondrashov 2014). To further evaluate whether asymmetrical evolution may be related to positive selection, we tested for positive selection using codeML (see Material and Methods for details). We detected positive selection in 388 genes distributed in 316 duplications (**Table S3, S4**), which supports that positive selection contributed to the asymmetrical acceleration of a substantial fraction of duplicates (at least ~37%). In addition, in most duplications, only one duplicate was under positive selection, with some exceptions where both duplicates showed signs of selection (28 and 44 for “strict duplicates” and “expansions” respectively, **Table S4**). Interestingly, post-duplication branches under positive selection have significantly different (and faster) rates than both the post-duplication branch without positive selection and the pre-duplication branch in both subsets (Fig. 3C), which is in agreement with the lower levels of identity detected for branches under positive selection (**Fig. S1B**, yellow boxplot). It is worth noting that our estimate of positive selection cases may be conservative due to the strict filtering applied and the inherent difficulty of detecting positive selection since this often acts during short periods of evolutionary time (Zhang 2005; Pegueroles et al. 2013; Pich I Roselló and Kondrashov 2014). However, a recent paper showed that the branch-site test (BST) cannot distinguish which sequence patterns have been caused by positive selection or by the neutral fixation of non-synonymous multinucleotide mutations (MNMs) (Venkat et al. 2018). For this reason, we identified MNMs in our gene duplicates. To do so, first, we reconstructed the ancestral sequence using codeML (see Material and Methods for details) and second, we compared the derived and ancestral sequences codon by codon and counted the number of changes. Those codons having more than one change were considered as MNMs. **Fig. S3** shows the percentage of MNMs per sequence for the duplicated genes, classified as having or not signals of positive selection. We observed that genes under positive selection are enriched in MNMs in both subsets. Therefore, we cannot rule out the possibility that some genes under adaptation according to the BST are actually due to the presence of MNMs.

**Figure 3:**
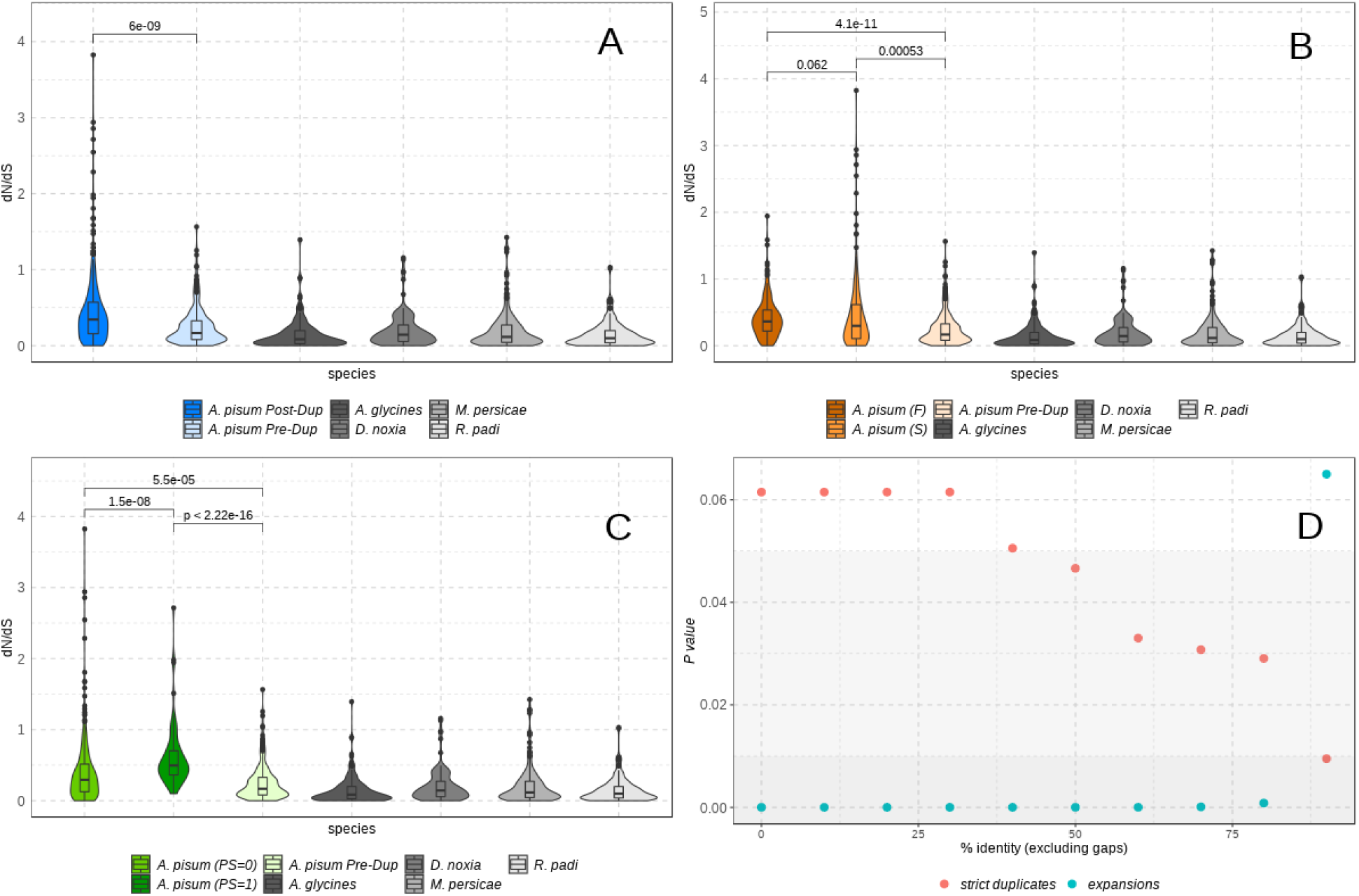
Evolutionary rate (dN/dS) for the selected duplications after filtering duplicates with dS >2 and dS <0.01 in any of the genes (201 strict duplicates remained). *A. pisum* genes are coloured while single-copy orthologs are shown in grey scale. **A:** *A. pisum* pre-duplication (Pre-Dup) and post-duplication (Post-Dup) branches are shown. **B:** *A. pisum* post-duplication branches were classified as Fast (F) or Slow (S) according to dS. **C:** *A. pisum* post-duplication branches were classified as having (PS=1) or not (PS=0) signals of positive selection. **D:** P-values comparing Fast (F) or Slow (S) copies from duplicates binned according to their percentage of identity (increasing by 10 the percentage of identity between bins). Background have been coloured according to the p-value: dark grey for p-value <0.01, light grey for p-value >0.01 and <0.05 and, white for p-value >0.05. P-values for all plots were estimated using wilcox.test function from R.

To evaluate the impact of positive selection in pairs of gene duplicates we discarded those duplicates with dS>2 or dS<0.01 in any of the duplicated genes (Fig. 4A, **Fig. S4**). The fraction of duplicates under positive selection is higher for the fast paralogs as compared to their slow counterparts (Fig. 4A, **Fig. S4**) which supports that the asymmetrical increase in rates may be due to adaptive selection, at least in a fraction of the duplications analysed, especially in the “strict duplicates” subset. We also observed that the fast post-duplication copies tend to have shorter sequence lengths in the “strict duplicates” subset (Fig. 4B, not in the “expansions” subset as shown in **Fig. S4B**). The evolutionary rates of the fast and slow evolving copies is quite similar independently of being consecutively positioned in the genome or not, at least in the “strict duplicates” set (Fig. 4C). It is worth noting that cases of positive selection are higher in the “expansions” subset, and positive selection is more prone to be wrongly assigned to this subset due to its complexity.

**Figure 4:**
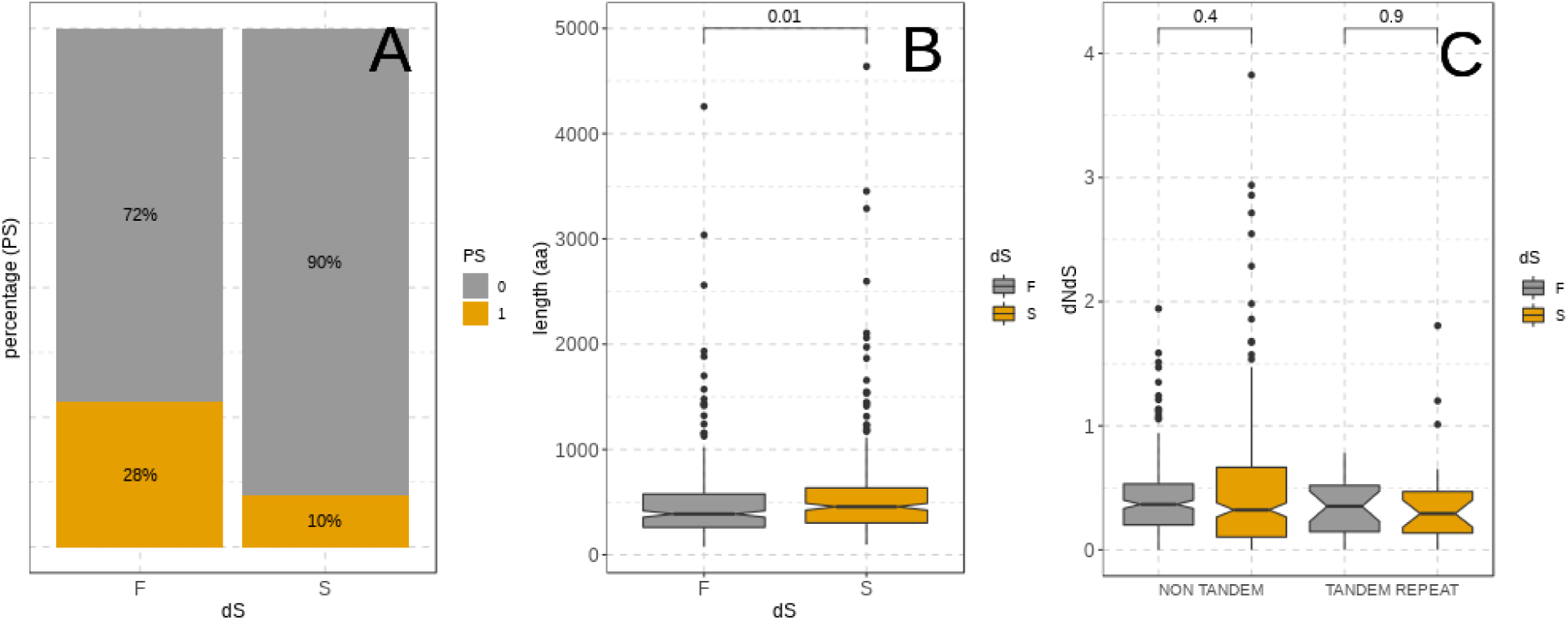
*A. pisum* genes from the selected duplications classified as Fast (F) or Slow (S) according to dS, after filtering duplications with dS >2 and dS <0.01 (201 strict duplicates remained, see Material and Methods for details). **A:** percentage of genes under positive selection (PS=1 in ochre) or with no signal of positive selection (PS=0 in grey); **B:** cDNA length (in aa); **C:** dN/dS after classifying duplicates according to their relative location (*i.e.* tandem and non-tandem duplicates). P-values were estimated using wilcox.test function from R.

We further evaluated the presence of selective pressures on *A. pisum* genome by estimating Tajima’s D for all genes annotated. We observed that overall values are negative, which implies an excess of rare alleles, which may be due to purifying selection or a recent population expansion after a bottleneck. When splitting the genome into “strict duplicates”, “expansions” and the rest of the genes (which are mostly not duplicated) we observed that “expansions” have significantly higher values than both “strict duplicates” and the rest of the genes, which may imply that the selective pressure to constrain “expansions” is lower (**Fig. S5**). However, no significant differences were found when comparing Tajima’s D for the fast and slow evolving copies in none of the subsets, indicating that overall both copies may underwent the same strength of purifying selection. Importantly, we observed that exons have significantly lower values compared to introns. Thus, negative Tajima’s D are more likely to be driven by purifying selection rather than a recent population expansion (**Fig. S6**), despite changes in population size are expected in a species that reproduces by cyclical parthenogenesis, such as the *A. pisum*. We also investigated codon usage index in gene duplicates, since departures from optimal codons in one of the copies would suggest relaxed purifying selection. Most of the duplicates (62 %) have an optimal codon usage, while in 32 % of the cases both copies have an non-optimal codon usage. Thus, our data set has a few cases in which only one copy departs from optimal codon usage. However, we explored whether for pairs of duplicates in which one of the copies was under positive selection was not adapted while its paralog was. Out of the 244 duplicates in which one gene is under positive selection only 9 pairs had the positively selected copy with a low codon adaptation index. A similar number (8 pairs) was found for pairs where the selected gene was the one better adapted to the codon usage. Thus, the set of recent duplicates does not seem to have a differential relaxation of purifying selection among copies. Altogether, these observations point that, at least in the “strict duplicates” subset, neofunctionalization fueled by positive selection seems the most likely scenario to explain the observed evolution patterns of recent duplicates in *A. pisum*.

To know the putative functions of the duplicated and positively selected genes, we tested whether the resulting paralogs were enriched in any particular functions through GO enrichment analyses. We explored enrichment only in strict duplicates as described above, which accounts for 72% of all duplicates. Similar results were found when the list of strict duplicates under selection was compared to all the genome or to the complementary portion of the genome (*i.e.*, all genes but the pairs of strict duplicates where at least one gene is under positive selection), including neurotransmitter metabolism, neural retina development, biosynthesis and metabolism of glutamate, quinone and ammonia (**Fig. S7**), suggesting that neofunctionalization may be affecting these functions. Strict duplicates under positive selection did not result in any functions enriched when compared to all strict duplicates.

A first example of a gene positively selected that may have undergone neofunctionalization is the gene encoding the protein *maelstrom 2* (UniProtKB - B3MZY6 MAEL2_DROAN), that in *Drosophila ananassae* has been predicted to play a central role during oogenesis by repressing transposable elements and preventing their mobilization, essential for maintaining the germline integrity (Sato et al. 2011). It is also the case of the genes encoding for *glutamine synthetase 2* (UniProtKB J9JML2_ACYPI) and other genes involved in glutamate metabolism such as the genes encoding aspartate aminotransferase 2 (UniProtKB J9JIS1_ACYPI), glutamate dehydrogenase (UniProtKB J9KB74_ACYPI) and glutamate decarboxylase (UniProtKB DCE_DROME). These genes belong to the same pathway of incorporation of ammonium nitrogen into glutamate cycle to assimilate ammonia into glutamate: those genes have been shown to be upregulated in bacteriocytes in *A. pisum*, which function as specialized symbiont-bearing organs of amino acid production (Hansen and Moran 2011). Therefore, positive selection and neofunctionalization may have facilitated host–symbiont cooperation in the production of amino acids between the pea aphid and *Buchnera*, such as for the amino-acid transporters as shown by Duncan et al. (2016). Overall, these two examples illustrate how key biological functions (oo- and morphogenesis and host-symbiont cooperation) might have been reshaped through duplication followed by neofunctionalization in the pea aphid.

### 2. Tissue divergence patterns in duplicated genes range from low to high

We have shown that recent *A. pisum* duplicates have different evolutionary rates and that a substantial fraction of them are evolving under positive selective pressure. The different behaviour of the two copies may result in differences in gene expression levels. To evaluate this hypothesis, we compiled RNA-Seq data from a total of 106 libraries grouped into 18 different conditions (see Materials and Methods). A principal component analysis showed that samples are clustered by tissues when considering all annotated genes in the *A. pisum* genome as well as when subtracting the genes of the “strict duplicates” and “expansions” subsets (**Fig. S8**). To compare gene expression profiles across tissues we profiled the expression of each selected gene, being 0 for not-expressed and 1 for expressed (**Table S5**, see Material and Methods for details). Interestingly, we observed that 329 duplicates (*i.e.* ~39% of the 843 selected duplicates, 212 of them “strict duplicates” and 117 “expansions”) showed differences in their tissue expression pattern in at least one tissue. When considering their location, we observed that corresponds to 40.4% of the 705 DNA-based pairs, 36.4% of the 44 retrocopies and 29.8% of the 94 tandem duplicates. Thus, most of tandem duplicates (~70%) tend to have the same expression patterns, as expected. In order to measure the expression divergence between duplicates in the 18 conditions, we computed three different statistics using a binary profiling binning approach: hamming distance, tissue expression complementarity (TEC) distance and tissue divergence (dT, **Table S4**, see Material and Methods for details). The three methods show similar results in both subsets (“strict duplicates” and “expansions”), which is in agreement with the high correlation between them (**Fig. S9**). Overall, tissue divergence between duplicates is low, with mean values ranging from 0.09 to 0.20 and the median being 0 in the three methods, which was expected since ~71% of the duplicates have the same expression profile. As expected, when considering merely pairs with differences in the expression profile we obtained higher values (median values ranged from 0.33 to 0.20). Interestingly, the maximum value detected is 1 for the three methods, meaning that some pairs of duplicates have totally opposite expression patterns.

We also compared gene expression between copies estimated as transcripts per million (TPMs). It is worth noting that overall gene expression values for “expansions” is significantly lower than for “strict duplicates” (median values were 1.07e-04 and 2.43e-05 respectively, p-value=2.64e-07). This may reflect real biological differences such as higher tissue specificity or lower expression of genes that are part of large family expansions, although it may also be due to the difficulty to assign gene expression to a given copy in these complex expansions and/or to the likely higher amount of missanotations or pseudogenes in this subset, as discussed above. We compared gene expression across tissues, by computing pearson correlations and building linear models within gene duplicates (see Material and Methods for details). If the two copies have similar expression patterns across tissues we should expect high pearson correlations and r squared values. Overall, our findings are in line with the binning approach, since both pearson correlations and r squared values are high (0.91 and 0.82 respectively for “strict duplications” and 0.77 and 0.60 for “expansions” (**Table S6**). In addition, the subset of differentially expressed genes obtained in the binning analysis is enriched in differentially expressed genes according to our models (Fig. 5, **Fig. S10**). Thus, we can conclude that the two subsets tend to have similar expression patterns in the two copies, despite the fact that there are some interesting differences as we will discuss below.

**Figure 5:**
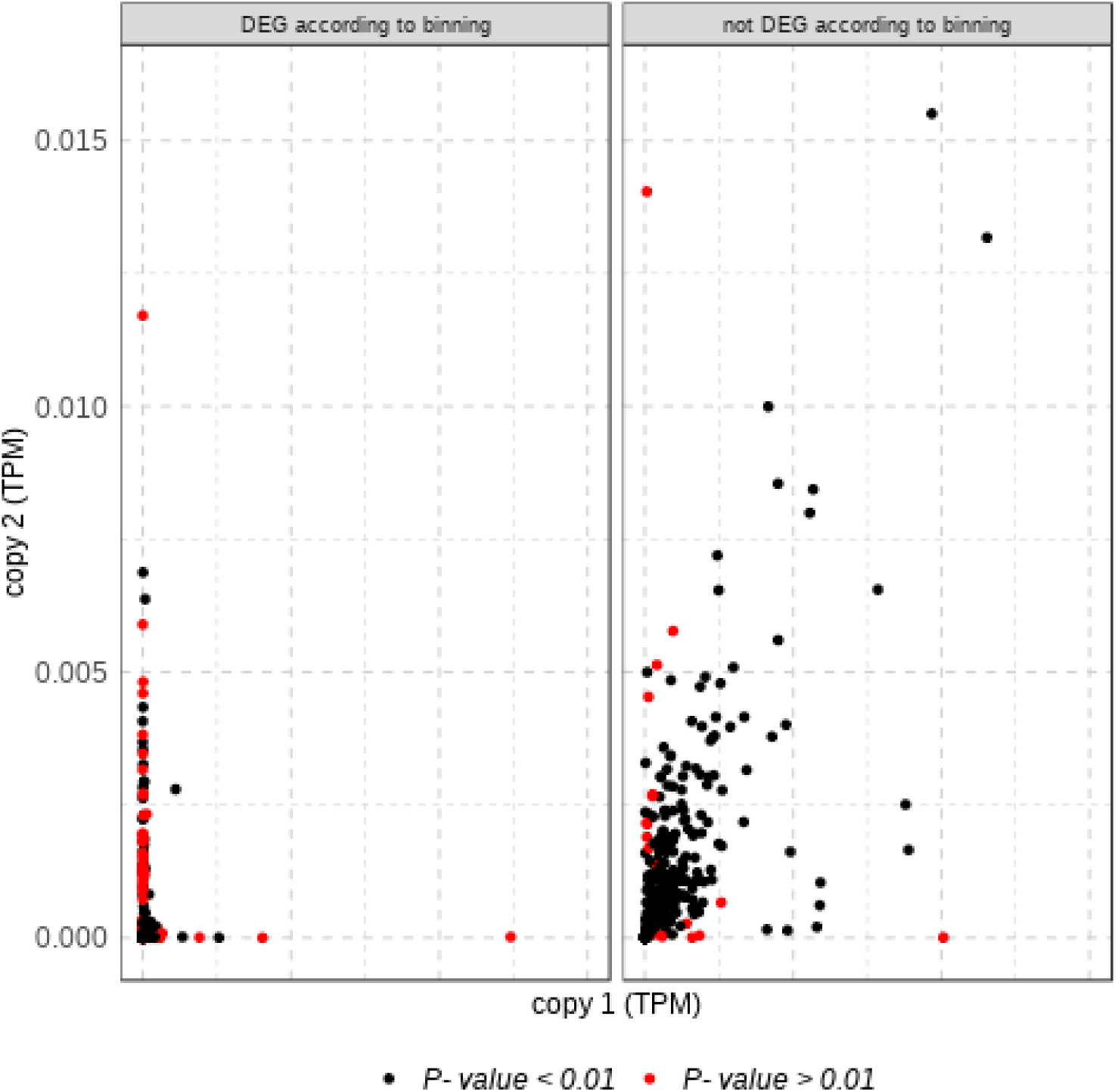
Scatterplot showing median gene expression between pairs of duplicates for the “strict duplications” subset. Colors correspond to the p-values of a Pearson correlation (see Material and Methods for details).

### 3. Positive selection may modulate differences in gene expression

Positive selection might be correlated with sub- or neofunctionalization by acquiring a new expression profile. To test this hypothesis, we compared the tissue expression patterns between gene duplicates. We found different expression patterns in 50% of pairs with two copies under selection (5 out of 14 “strict duplicates” and 13 out of 22 “expansions”), 36.9% of pairs with only one copy under selection (52 out of 173 “strict duplicates” and 36 out of 71 “expansions”) and 39.6% of pairs with no copy under selection (155 out of 419 “strict duplicates” and 68 out of 144 “expansions”). This suggests that positive selection plays a role in gene transcription regulation but other factors are also involved, since in the absence of positive selection, differences in gene expression were also detected. When focusing on duplications that have different expression patterns in at least one of the studied conditions, we observed that tissue expression divergence levels were similar for duplications having or not copies under selection in both subsets (“strict duplicates” and “expansions”) as well as overall duplicates (**Fig. S11**). For the 244 duplications with positive selection in one copy, we quantified the cases in which a gene expression was gained or lost in any of the tissues considering the expression profile of the copy with absence of positive selection as background (**Table S5**). The number of losses was higher than that of gains in the “strict duplicates” subset (40 and 15 respectively), meaning that in most cases the gene expression profile of the copy under selection is reduced as compared to the non-selected copy. In other words, the selected copy is expressed in a subset of tissues at least in the “strict duplicates” subset, since the median number of tissues in which the non-selected and the selected copies are expressed is 11 and 8.5, respectively, in this set of duplicates. In the “expansions” subset the amount of gains and losses is quite similar (21 and 19 respectively), as well as the median number of tissues in which the non-selected and the selected copies are expressed (9 and 10 respectively). The pattern observed in the “strict duplicates” subset is consistent with a specialization scenario, in which one copy is expressed in all (or most) tissues but at least one copy is not. This scenario, which can be considered a particular case of subfunctionalization, has been proposed to be the main fate after whole genome duplication (Marlétaz et al. 2018) and may influence the evolution of young duplicates (Huerta-Cepas, Dopazo, et al. 2011). In addition, the 15 cases in which the selected copy is expressed in at least a tissue in which the non-selected copy have no expression (gain cases) are candidates that may have undercome neofunctionalization after gene duplication (Table 1). From these 15 duplications, 9 showed similar predicted annotation between the pairs (duplications 288, 322, 397, 460, 489, 576, 633, 795, 840). The 6 other duplications have made of pairs with different predicted annotations. It is worth noting that for the 15 duplications (i.e., 30 genes), 16 are uncharacterized no predicted functions, 5 are annotated as zinc-finger putative proteins, and 3 are dynein-like proteins.

**Table 1:** Duplications in which the copy under positive selection (PS=1) is expressed in at least a tissue in which the non-selected copy (PS=0) have no expression (highlighted in dark grey). NA, no information on gene expression available. (see Material and Methods for further details about each condition type). Abbreviations: AM: Adult Males. AP: Adult Females Parthenogenetic. AO: Adult Female Oviparae. E0: Embryos Stage17. E1A: Embryos Stage18 Sex. E1K: Embryos Stage18 Asex. E2A: Embryos Stage19 Sex. E2K: Embryos Stage19 Asex. E3A: Embryos Stage20 Sex. E3K: Embryos Stage20 Asex. H: Head_Mix. HP: Head Adult Female Parthenogenetic. HR2: Head Larvae2. HR4: Head Larvae4. LP: Legs Adult Female Parthenogenetic. G: Gut. SG: Salivary glands. B: Bacteriocyte.

| Duplication name | Gene code | Putative function | P S | A M | A P | A O | E 0 | E 1A | E 1K | E 2A | E 2K | E 3A | E 3K | H | H P | H R2 | H R4 | L P | G | SG | B |
| --- | --- | --- | --- | --- | --- | --- | --- | --- | --- | --- | --- | --- | --- | --- | --- | --- | --- | --- | --- | --- | --- |
| duplication_159 | LOC103311559 | Uncharacterized | 0 | 0 | 0 | 0 | 0 | 0 | 0 | 0 | 0 | 0 | 0 | NA | 0 | 0 | 0 | 0 | 0 | NA | 0 |
| duplication_159 | LOC100166252 | PiggyBac transposable element-derived protein 4-like | 1 | NA | NA | NA | NA | NA | NA | NA | NA | 1 | 1 | NA | NA | 0 | 0 | 0 | 1 | NA | NA |
| duplication_213 | LOC100570324 | Uncharacterized | 0 | 1 | 0 | NA | NA | 1 | 1 | NA | 1 | NA | NA | 0 | 1 | NA | 0 | 0 | NA | NA | NA |
| duplication_213 | LOC100574933 | PAX-interacting protein 1-like | 1 | 1 | 1 | 1 | 1 | 1 | 1 | 1 | 1 | 1 | 1 | NA | NA | 1 | 1 | 1 | NA | 1 | 1 |
| duplication_288 | LOC100572588 | Uncharacterized | 0 | 1 | 1 | 1 | 1 | 1 | 1 | 1 | 1 | 1 | 1 | 1 | 1 | 0 | 0 | 1 | 1 | 1 | 1 |
| duplication_288 | LOC103308356 | Uncharacterized | 1 | 0 | 1 | 1 | 1 | 1 | 1 | 1 | 1 | 1 | 1 | 1 | 1 | 1 | 1 | NA | 1 | 1 | 1 |
| duplication_322 | LOC107883251 | Uncharacterized | 0 | 0 | NA | NA | 1 | NA | 1 | 1 | 1 | 1 | 1 | 1 | NA | 1 | 1 | NA | 1 | 1 | 1 |
| duplication_322 | LOC107883068 | Uncharacterized | 1 | 1 | 1 | NA | 1 | 1 | 1 | 1 | 1 | 1 | NA | 1 | 1 | 1 | 1 | 1 | 1 | 1 | 1 |
| duplication_340 | LOC103310866 | Uncharacterized RING finger protein C32D5.10-like | 0 | 0 | 0 | 0 | NA | 0 | 0 | 0 | 0 | 0 | 0 | 0 | 0 | 0 | 0 | 0 | NA | 0 | 0 |
| duplication_340 | LOC100571229 | E3 ubiquitin-protein ligase Topors-like | 1 | 1 | 1 | 1 | 1 | 1 | 1 | 1 | 1 | 1 | 1 | 0 | NA | NA | 0 | 1 | NA | NA | 1 |
| duplication_397 | LOC100162340 | Dynein heavy chain 1, axonemal | 0 | 1 | 0 | 0 | 0 | 0 | 0 | 0 | NA | 0 | 0 | 0 | 0 | 0 | NA | 0 | NA | 0 | 0 |
| duplication_397 | LOC107882216 | Dynein heavy chain 1, axonemal-like | 1 | 0 | NA | 1 | 0 | 0 | 0 | NA | 0 | NA | 0 | NA | 1 | 0 | NA | 0 | NA | NA | NA |
| duplication_4 | LOC100568916 | Uncharacterized | 0 | 1 | NA | 1 | 1 | 1 | 1 | 1 | 1 | 1 | 1 | NA | NA | 0 | 0 | NA | NA | NA | NA |
| duplication_4 | LOC100160128 | 26S proteasome non-ATPase regulatory subunit 12-like | 1 | 1 | 1 | 1 | 1 | 1 | 1 | 1 | 1 | 1 | 1 | NA | 1 | 1 | 1 | 1 | NA | NA | 1 |
| duplication_460 | LOC107882168 | Zinc finger protein 134-like | 0 | 0 | 0 | NA | 0 | NA | NA | 0 | 0 | NA | NA | 0 | 0 | 0 | 0 | 0 | NA | NA | 0 |
| duplication_460 | LOC103310004 | Zinc finger protein 134-like | 1 | 1 | 0 | 1 | NA | NA | 0 | 0 | NA | 0 | 0 | 0 | 0 | 0 | 0 | 0 | NA | NA | 0 |
| duplication_489 | LOC107882711 | Uncharacterized | 0 | NA | 1 | 1 | 1 | 1 | 1 | 1 | 1 | 1 | 1 | NA | 1 | 1 | 1 | 0 | 1 | NA | 1 |
| duplication_489 | LOC100162929 | Uncharacterized | 1 | 1 | 1 | 1 | 1 | 1 | 1 | 1 | 1 | 1 | 1 | 1 | 1 | 1 | 1 | 1 | 1 | 1 | 1 |
| duplication_576 | LOC100569858 | Uncharacterized | 0 | 1 | 0 | 0 | NA | NA | 0 | 0 | 0 | 0 | 0 | NA | 0 | 0 | 0 | 0 | NA | NA | 0 |
| duplication_576 | LOC100574180 | Uncharacterized | 1 | 1 | 0 | 0 | NA | NA | 0 | NA | NA | 0 | NA | 0 | NA | 1 | NA | 0 | 1 | 1 | 0 |
| duplication_633 | LOC103309550 | Zinc finger MYM-type protein 1-like | 0 | 0 | 0 | 0 | 0 | 0 | 0 | 0 | 0 | 0 | 0 | NA | 0 | 0 | 0 | 0 | NA | NA | 0 |
| duplication_633 | LOC103307955 | Zinc finger MYM-type protein 1-like | 1 | 0 | NA | NA | 0 | NA | 0 | 0 | NA | 0 | 0 | NA | NA | NA | 0 | 0 | NA | NA | 1 |
| duplication_663 | LOC107884962 | Uncharacterized | 0 | 0 | 0 | 0 | 1 | NA | 1 | NA | NA | NA | 1 | NA | 0 | 0 | 0 | 0 | 1 | 1 | NA |
| duplication_663 | LOC100570263 | Uncharacterized | 1 | NA | 1 | 1 | 1 | 1 | 1 | 1 | 1 | 1 | 1 | NA | 1 | NA | 0 | 1 | NA | 1 | 1 |
| duplication_693 | LOC100165085 | Uncharacterized | 0 | 1 | 0 | 0 | NA | 0 | NA | 0 | NA | 0 | 0 | NA | 0 | NA | NA | NA | NA | NA | 1 |
| duplication_693 | LOC100165046 | Dynein heavy chain 7, axonemal-like | 1 | NA | 1 | 0 | 1 | 1 | 1 | 1 | 1 | 1 | 1 | 1 | 1 | 1 | 1 | 1 | 1 | 1 | 1 |
| duplication_795 | LOC107882727 | Sialin-like | 0 | 0 | NA | 0 | 0 | 0 | 0 | 0 | 0 | 0 | 0 | NA | 1 | 1 | 1 | 1 | NA | NA | NA |
| duplication_795 | LOC100164217 | Putative inorganic phosphate cotransporter | 1 | 1 | 1 | 1 | NA | 1 | 1 | 1 | 1 | 1 | 1 | 1 | 1 | 1 | 1 | 1 | 1 | 1 | 1 |
| duplication_840 | LOC100571978 | Uncharacterized | 0 | 1 | 0 | 0 | 0 | 0 | 0 | 0 | 0 | 0 | 0 | 0 | 0 | 0 | 0 | 0 | 0 | NA | 0 |
| duplication_840 | LOC107884413 | Uncharacterized | 1 | 0 | 0 | 0 | 0 | NA | NA | 0 | NA | NA | 0 | NA | NA | 1 | 1 | 0 | NA | NA | 0 |

### 4. Chromatin accessibility is altered in young duplicated genes but does not correlate to gene expression

RNA-Seq and FAIRE-Seq data were analysed together for each of the predicted genes in the *A. pisum* genome (**Table S7**). Faire-seq is a molecular technique that allows to detect nucleosome-depleted regions of the genome, which are usually found in open chromatin. We validated the overall correlation of transcription and chromatin accessibility (**Figure S12** for embryos and Richard et al. 2017, Figure 5 for whole-body males and females), ensuring the quality of the datasets. In order to explore the correlation between transcriptional status and chromatin accessibility, genes were classified in four categories depending on their expression and TSS log2 (FAIRE/Input) values (see Methods): (i) open and expressed, (ii) open and not expressed, (iii) closed and expressed, and (iv) closed and not expressed. Concerning all genes, we found that most (8,870 genes; 64%) belonged to the category “closed and expressed”, which could potentially be due to the threshold applied to define “open genes” and to the higher sensitivity of RNA-Seq compared to FAIRE-Seq. From the remaining, 18% of the genes were “open and expressed”, and the other 18% “closed and not expressed” (2,497 and 2,491 genes, respectively). The “open and not expressed” category was barely represented in the dataset (0.3% of the assigned genes). This result reflects the quality of the FAIRE-Seq data processing since it is expected that “open and not expressed” genes are virtually absent as they will violate the common rules of gene transcription, therefore corresponding to false positives. GO term enrichment analysis for each of the four categories revealed that genes “open and expressed” were enriched in transcription factor activity (GO:0003700, molecular function) and sequence-specific DNA binding (GO:0043565, molecular function). Genes “closed and not expressed” were enriched in nucleic acid binding (GO:0003676, molecular function). The two remaining categories were not significantly enriched in any functions, despite the high number of genes in the “closed and expressed” category: this highlights the biological relevance of gene classes displaying coordinated expression and TSS accessibility in this dataset.

Regarding the set of young gene duplicates, we found that 26% of them had different chromatin states in each paralog (173 out of 666 duplicates, **Table S8**). If we divide young duplicates into “strict duplicates” and “expansions” we observe similar patterns (26% and 27% respectively). Therefore, the duplicates, no matter whether a simple pair of a more complex duplication schema, potentially have different expression patterns. To test this hypothesis, we searched for duplicates where each paralog belonged to a different category (*i.e.*, categories (i) to (iv) combining chromatin state and expression pattern, as described above) for the embryos and adult morphs conditions for which FAIRE-Seq data were available (**Table S8**). Seventy three pairs of duplicated genes did belong to different categories. From those, in 54 pairs both duplicates were expressed, with one copy being open and the other closed. In 19 pairs, both duplicates were closed but one gene was expressed and the other one not.

Our results indicate that the number of genes with closed chromatin (n=11,322) was higher than for open chromatin (n=2,536). This reduces overall the possible correlation of genes expressed and accessible at the same time, consequently hampering any gene-by-gene comparisons such as in the case of duplicated genes. Our results are in line with those of the integration of RNA-Seq and ATAC-Seq (similar to FAIRE-Seq) performed in a study (Ackermann et al. 2016), who discussed that the poor correlation between RNA-Seq and ATAC-Seq data in human tissues may be due to gene activation depending on multiple regulatory regions, possibly being located far from the gene locus itself. Indeed, FAIRE-Seq in whole body individuals or embryos is far less precise than at the level of cells or tissues, thus making the correlation between expression and accessibility even trickier. Also, since FAIRE-Seq only allows to test for *cis*-regulatory interactions, it may be hypothesized that most of the genes may be trans-regulated, which was impossible to determine with the data at hand considering the non-completeness of the pea aphid genome assembly. Moreover, our analysis was centered on putative TSS which have not been validated experimentally, for instance by CAGE-Seq (Takahashi et al. 2012) or NET-Seq (Churchman and Weissman 2012) or MAPCap (Bhardwaj et al. 2019). Nevertheless, we identified that the chromatin accessibility of the TSS of duplicated genes was different between the pairs in more than half of the cases. This shows that the chromatin states of promoters of simple duplicated genes can evolve independently in each pair. This could correspond to the differential chromatin states recently identified in new genes in nematodes (Werner et al. 2018), since in each pair, one gene is more recent than the other.

## Conclusions

Our study shows that recent gene duplicates in *A. pisum* evolved asymmetrically, with one more conserved and one more divergent paralogous copy. The conserved copy was likely maintained mainly through purifying selection pressures and hardly ever under the effect of positive selection. The divergent copy was usually positively-selected and showed a faster evolutionary rate. Altogether, these results suggest that neofunctionalization may be one of the driving forces affecting young gene duplicates in *A. pisum.* In addition, genes under positive selection were putatively related to a large and diverse number of functions, indicating that neofunctionalization has a broad impact in multiple functions of the pea aphid biology. Remarkably, neofunctionalization may also be involved in symbiosis functioning, facilitating host-symbiont cooperation between *A. pisum* and *Buchnera*. Conclusions are strong for strict duplicates (i.e. genes that merely duplicated once) but more complex scenarios may have driven the evolution of large expansions, which remain difficult to analyze due to confounding factors, such as possible missannotations and pseudogenes. Concerning the expression patterns of the duplicated genes, we observed that more than one third of the duplicates showed different expression patterns, with some of them under adaptive selection. This suggests that positive selection might not be the main or the only factor driving these differences in gene expression. For those duplicates with signals of positive selection, we found that a loss of function in a specific tissue is the most likely outcome, consistent with a scenario of tissue specialization and/or subfunctionalization. In contrast, we also found examples of genes under positive selection that gained their function in some tissues, compatible with a scenario of neofunctionalization.

Lastly, we did not find a relationship between chromatin accessibility and gene expression, which may potentially be explained by technical issues such as a limited prediction of TSS in the pea aphid genome coupled to the inherent low signal over background ratio of FAIRE-Seq data (Ackermann et al. 2016). Moreover, although this discrepancy may be due to the different sensitivity of both RNA-Seq and FAIRE-Seq, it may also reflect a pervasive level of *trans* regulation in the pea aphid genome (as seen in humans (Ackermann et al. 2016)). Nevertheless, we showed that more than half of the young duplicated genes selected had different chromatin states. This indicates that FAIRE-Seq technique is sensitive to differences in chromatin dynamics even in recent gene duplicates.

Altogether, our results indicate that gene duplication provided an arena of genetic novelty to reshape the genome of the pea aphid through positive selection, neofunctionalization and tissue-specific expression in young duplicated species-specific genes. The relationships between these evolutionary scenarios are complex and difficult to disentangle. We emphasize that phylogenomic-centered studies are therefore most needed to further understand genome evolution in nonmodel organisms.

## Material and methods

### 1. Identification and selection of duplications in the pea aphid genome

The phylome (*i.e.*, the complete collection of phylogenetic trees for each gene in its genome) of *A. pisum* Mordvilko, 1914 was reconstructed in the context of Hemiptera evolution. In addition to this species, belonging to the suborder Sternorrhyncha and to the family Aphididae and tribe Macrosiphini, we selected representatives of several hemipterans based on phylogenetic position and availability of a fully-sequenced genome: *Diaphorina citri* Kuwayama, 1908 (Sternorrhyncha, Psylloidea), *Bemisia tabaci* (Gennadius, 1889) (Sternorrhyncha, Aleyrodoidea), *Daktulosphaira vitifoliae* (Fitch, 1855) (Sternorrhyncha, Phylloxeridae), *Cinara cedri* (Curtis, 1835) (Sternorrhyncha, Aphidoidea), *Diuraphis noxia* (Kurdjumov, 1913) (Sternorrhyncha, Aphidoidea), *Aphis glycines* (Matsumara, 1917) (Sternorrhyncha, Aphidoidea), *Myzus persica*e (Sulzer, 1776) (Sternorrhyncha, Aphidoidea) and *Rhopalosiphum padi* (Stal, Linnaeus, 1758) (Aphidinae, Aphidini) (Fig. 1). Genome versions and number of predicted proteins are indicated in **Table S1**.

Phylomes were reconstructed using the PhylomeDB pipeline (Huerta-Cepas, Capella-Gutierrez, et al. 2011). For each protein encoded in the *A. pisum* genome, a BLAST search was performed against the custom proteome database built from the genomes listed above. Results were filtered using an e-value of 1e-05 and a minimum overlapping region of 0.5. Multiple sequence alignments were reconstructed in both directions using three different programs (MUSCLE v3.8 (Edgar 2004), MAFFT v6.712b (Katoh 2005), and Kalign (Lassmann and Sonnhammer 2005)) and combined using M-COFFEE (Wallace et al. 2006). A trimming step was performed using trimAl v1.3 (Capella-Gutierrez et al. 2009), consistency-score cutoff = 0.1667 and gap-score cutoff = 0.9). Following model selection, the best model in terms of likelihood as selected by the Akaike Information Criterion (AIC) was chosen for tree reconstruction. Phylogenetic trees were inferred using PhyML v3.0 (Guindon et al. 2010). Four rate categories were used and invariant positions were inferred from the data. Branch support was computed using an aLRT (approximate likelihood ratio test) based on a chi-square distribution. Resulting trees and alignments are stored in PhylomeDB 4.0 (Huerta-Cepas et al. 2014), http://phylomedb.org). The phylomeID is 441 (http://phylomedb.org/phylome_441).

A species-overlap algorithm, as implemented in ETE v3 (Jaime Huerta-Cepas et al. 2010) was used to infer orthology and paralogy relationships from the phylogenetic trees reconstructed in the phylome. The algorithm scans the tree and calls speciation or duplication events at internal nodes based on the presence of common species at both daughter partitions defined by the node. Gene gains and losses were calculated on this basis. Duplication ratios per node were calculated by dividing the number of duplications observed in each node by the total number of gene trees containing that node: theoretically, a value of 0 would indicate no duplication, a value of 1 an average of one duplication per gene in the genome, and >1 multiple duplications per gene and node.

To build the species tree, one-to-one orthologs present in all species were selected, resulting in a final alignment with 1,047 genes and 635,610 amino acid positions after concatenation. To ensure a congruent phylogenetic hypothesis under different models, a series of approaches were followed to infer the species tree. First, an ML tree was reconstructed with PhyML under the best selected model of amino acid evolution (LG, Le et al. (2008). Second, a supertree was reconstructed using DupTree (Wehe et al. 2008) based on all the trees reconstructed in the phylome. Both phylogenies were congruent (Fig. 1).

### 2. Detection and selection of gene duplications

For each gene tree, we first selected with ETE v3 (Jaime Huerta-Cepas et al. 2010) the nodes that exclusively contained multiple *A. pisum* sequences. These were considered species specific duplications in *A. pisum*. Overlapping species-specific duplications were fused when they shared more than 50% of their members. Trees were then scanned for the presence of pairs of duplicates in *A. pisum* whose duplication node was highly supported (aLRT > 0.95) and which had at least two single copy orthologs. Note that the selected pairs of duplicates are not limited to genes that duplicated just once, some of them belong to a (larger) expansion in which case the chosen pairs were always at the tips of the tree. Species-specific duplicated genes and selected orthologs were grouped and used to build a second ML tree. The purpose of this tree was to ensure that the resulting topology still contained the species specific duplication. Pairs of duplicates with incongruent CDS annotation or unsatisfactory topology. This resulted in a final number of scrutinized duplications of 843. For each duplication, we obtained multiple protein sequence alignments with PASTA v1.8.3 (Mirarab et al. 2015) and computed a gene tree (using the tree-estimator RAxML option) that was used for codeml analyses (see below). We finally back-translated protein multiple sequence alignments into nucleotidic with trimAl (using -phylip_paml - nogaps -backtran options). We also estimated median identity for each protein sequence in the alignment using trimAl v1.3 (-sident option after omitting gaps; Capella-Gutierrez et al. 2009). Gene conversion was estimated from back-translated sequences using GENECONV software (Sawyer 1989), by considering fragments with evidence of a gene conversion event between the ancestors of two *A. pisum* sequences that remained significant after multiple-comparison correction.

### 3. Evolutionary rates and additional filtering

We estimated the number of synonymous substitutions per synonymous site (dS), the number of non-synonymous substitutions per non-synonymous site (dN) and dN/dS ratio using the “free ratio branch model” implemented in codeML from PAML v. 4.9 (Yang 2007), using model

= 1, CodonFreq = 3, Nsites = 0 as options. This software allows to estimate dS, dN and dN/dS for each internal and terminal branch of a given tree and also to reconstruct the ancestral sequence before the duplication occurred. Analyses were computed for the 1020 selected duplications, which contained a specific duplication in *A. pisum* and at least two single-copy orthologs for any of the other eight species included in the phylome. We noticed that dS for the three more distant species was much higher (the percentage of sequences with dS>2 was 82.1%, 86.2% and 63.9% in *D. citri*, *B. tabaci* and *D. vitifoliae*, respectively) than in the closely related species (16.5%, 6.4%, 5.8%, 3.3%, 4.4% and 0.7% in *C. cedri*, *A. glycines*, *R. padi*, *D. noxia*, *M. persicae* and *A. pisum* respectively). Since such large dS values may indicate problems in the orthology identification we discarded duplications with only single-copy orthologs in the three most distant species. A total of 843 duplications remained after this filtering.

### 4. Age of the selected duplications and classification into fast and slow copies

The relative age of the selected duplications was calculated using the number of synonymous substitutions per synonymous site (dS) as a proxy. From the total of 5,589 genes from six species in 843 duplications we filtered out genes with dS >2 (which may indicate problems in the orthology identification, 242 genes) and dS < 0.01 (which may lead to high dN/dS ratios with no biological sense, 908 genes). We also used the dS estimates to classify the two copies of each selected duplication into fast and slow, by comparing their dS values, the copy with the lowest dS value being classified as slow and the other as fast.

### 5. Classification of gene duplications

Gene duplicates can be divided into two groups: strict duplicates (606), including *A. pisum* genes that derive from a recent common ancestor and duplicated specifically in this species only once (*i.e.*, genes with only two copies) and expansions (237), including *A. pisum* genes that also derive from a recent common ancestor but duplicated multiple times (*i.e.*, genes with more than two copies). In addition, duplicates were further classified as tandem (defined as duplicates with no genes in between; 94 in total), located in the same or different contig (dispersed duplicates, 100 and 605 respectively) and retrotransposed (defined as dispersed duplicates in which one copy lacks introns; 44 in total).

### 6. Selection tests

We tested for positive selection using the “branch-site” test 2 implemented in codeML from PAML v.4.9 (Yang 2007). We compared the null hypothesis where dN/dS is fixed in all branches (model = 2, NSsites = 2, fix_omega = 1, omega = 1) and the alternative hypothesis where the branch that is being tested for positive selection may include codons evolving at dN/dS>1 (model = 2, NSsites = 2, fix_omega = 0, omega = 1.5). The two models were compared using a likelihood ratio test (LRT) and p-values were adjusted for multiple comparisons using the Holm, Hochberg, SidakSS, SidakSD, BH and BY methods using multtest package for R. We considered that a given gene is under selection if any of the adjusted p-values computed using the different methods was < 0.01.

To compute Tajima’s D for each gene we mapped the reads with bwa and generated a pileup with samtools mpileup (Etherington et al. 2015). From the pileup file, we run the script subsample-pileup.pl from Popoolation (Kofler et al. 2011), with the option --target-coverage 15 -- max-coverage 150 --method withoutreplace. Then, Tajima’s D was calculated using the script "Variance-at-position.pl --measure D" from Popoolation, with a population size of 40, on a GTF file with all the protein coding genes in the *A. pisum* genome.

The codon adaptation index for each gene of interest was estimated using CaiCAL (Puigbò et al. 2008). The Codon usage table for A. pisum was obtained from CoCoPUTs (Athey et al. 2017). CaiCAL was also used to calculate the expected CAI value based on a 1000 randomly created sequences. Genes with a CAI value above the eCAI value were considered as optimized.

### 7. Functional annotation and GO term enrichment analysis and visualization

To assign Gene Ontology (GO) terms to the genes in the pea aphid genome, GO terms based on orthology relationship were propagated with eggNOG-mapper (Huerta-Cepas et al. 2017). For that, we selected the eukaryotic eggNOG database (euNOG; Huerta-Cepas et al. 2019) and prioritised coverage (*i.e.*, GO terms were propagated if any type of orthologs to a gene in a genome were detected). See **Table S9** for the full annotation of the selected genes. Functional enrichment of the selected duplications was explored with FatiGO (Al-Shahrour et al. 2004). We tested enrichment against two different backgrounds: all the genome and the remaining genes in the genome (*i.e.*, non-expanded genes and non-positively selected ones, respectively). Sets of GO terms were summarized and visualized in REVIGO (Supek et al. 2011).

### 8. Tissue expression diverge between duplicates

Messenger RNA (mRNA) expression data was obtained from 106 different samples from the *A. pisum* LSR1 lineage (The International Aphid Genomics Consortium 2010). We obtained RNA-Seq libraries from 18 different conditions. Some of them were retrieved from the public databases and others newly generated for this study (**Table S2**). These were sequenced using Illumina technology as paired-end of 100 bp size, containing more than 25 million raw reads per library. Reads from all the RNA libraries were mapped on the version 2 of the pea aphid genome assembly (Acyr_2.0, ID NCBI: 246238) using STAR version 2.5.2a (Dobin et al. 2013) with the default parameters except the following parameters: outFilterMultimapNmax = 5, outFilterMismatchNmax = 3, alignIntronMin = 10, alignIntronMax = 50000 and alignMatesGapMax = 50000. The number of reads covering each gene prediction (NCBI Annotation release ID: 102) was then counted using FeatureCounts version 1.5.0-p3 (Liao et al. 2014) with the default parameters except the following parameters: -g gene -C -p -M --fraction. For each counting, RPKM calculation was performed using edgeR (Robinson et al. 2010; McCarthy et al. 2012) with gene.length = sum of exons size for each gene. TPM were calculated from RPKM using the equation TPM(i) = (FPKM(i) / sum (FPKM all transcripts)) * 10^6. Principal components analysis (PCA) was performed using prcomp function from R.

RNA-Seq values for each individual gene were divided into four quartiles. Each RNA-Seq experiment was processed independently. Replicates were then joined by collapsing the different values obtained in the different experiments of the same tissue. If more than 50% of the experiments placed the RNA-Seq data into the same, this bin was assigned to the overall tissue. On the other hand, if none of the bins had enough representation across experiments, no value was assigned (NA). Once each tissue was assigned a value, a profile was created for each individual gene. The profiles consisted of 0 and 1 in which 0 represented not-expressed and were values located in the lowest of the four bins. 1 represented expressed genes and consisted of values located in the other three bins. These expression profiles were used to calculate the tissue expression divergence between pairs of duplicates using three different methods: i) Normalized Hamming distance, which counts the number of differences between two profiles and divides it by the total number of considered tissues. A tissue is not considered when its value is NA for either gene. ii) Tissue expression complementarity (TEC) distance (Huerta-Cepas, Dopazo, et al. 2011), which compares the relative number of tissues in which only one set but not the other was expressed over the total number of tissues in which each gene is expressed. iii) Tissue expression divergence (dT) (Pegueroles et al. 2013), which subtracts tissues were one or two copies are expressed from tissues were the two copies are expressed divided by tissues were one or two copies are expressed. Values for the three distances range from 0 to 1, where 0 means no differences in gene expression between duplicates (in other words, the two copies tend to be expressed in the same tissues) and 1 means that the two copies have totally different expression patterns. In addition, for each replicate of a given tissue we computed the median expression value in TPM. We then rescaled the expression across the tissues using the rescale function from plyr package from R and subsequently we calculated the median expression value, the pearson correlation and its p-value, and the r-squared and the slope of a linear model using gene expression across the 18 tissues.

### 9. FAIRE-Seq data analysis

FAIRE-Seq data for samples for males and females adults was taken from Richard et al. (2017). FAIRE-Seq samples for embryos were newly generated for another, unpublished, study (Richard 2017). Subsequently to sequencing, FAIRE and Control reads were mapped using bowtie2 with default parameters (Langmead et al. 2009; Langmead and Salzberg 2012) on the pea aphid genome assembly Acyr_2.0 (ID NCBI: 246238, AphidBase: http://bipaa.genouest.org/is/aphidbase/acyrthosiphon_pisum/). Only uniquely mapped reads with a mapping quality over or equal to 30 in the phred scale were kept using SAMtools (Li et al. 2009), following the IDR recommendations (Li et al. 2011), https://sites.google.com/site/anshulkundaje/projects/idr#TOC-Latest-pipeline). MACS2 (Zhang et al. 2008) was used to perform the peak calling with the following parameters using control samples: --gsize 541675471 --nomodel --extsize 500 -p 0.05 --keep-dup all -f BEDPE, followed by Irreproducible Discovery Rate (IDR) analyses using a threshold of 0.01 for original replicates, of 0.02 for self-consistency replicates and of 0.0025 for pooled pseudoreplicates. Replicates consistency was then assessed using the IDR algorithm (Li et al. 2011) and the two most correlated FAIRE replicates out of the three in each condition were pooled in order to reduce the noise, as widely recommended for ChIP-Seq or ATAC-Seq data. Input-normalized FAIRE-Seq signals were calculated using deepTools2 *‘bamCompare’(Ramírez et al. 2016)* across the whole genome for each condition by calculating the average log2 (Pooled FAIRE/Input) in windows of 10 bp. Both Pooled FAIRE and Input read counts were normalized by sequencing depth using *‘--normalizeTo1x’*. Using deepTools2 *‘multiBigwigsummary’*, the average FAIRE signal was extracted 900 bp around the beginning of genes (450 bp in 5’ and 450 bp in 3’) in all samples. We then used a threshold of 1 for the average log2 (FAIRE/Input) to define genes whose TSS is open (above the threshold) or closed (below the threshold).

Embryos and adults RNA-Seq data were related to the FAIRE-Seq data for each condition and individual gene. According to the data, genes were classified in four categories: (i) open and expressed, (ii) open and not expressed, (iii) closed and expressed and (iv) closed and not expressed. For each gene, the percentage of tissues representing each category was calculated. If this average reached at least 75%, a single category was assigned to the gene. 13,858 genes were assigned to one of the categories (**Table S6**).

## Acknowledgements

RF was funded by a Juan de la Cierva-Incorporación Fellowship (Government of Spain, IJCI-2015-26627) and a Marie Skłodowska-Curie Fellowship (747607). TG acknowledges support from the Spanish Ministry of Economy, Industry, and Competitiveness (MEIC) for the EMBL partnership, and grants ‘Centro de Excelencia Severo Ochoa 2013-2017’ SEV-2012-0208, and BFU2015-67107 co-funded by European Regional Development Fund (ERDF); from the CERCA Programme / Generalitat de Catalunya; from the Catalan Research Agency (AGAUR) SGR857, and grant from the European Union’s Horizon 2020 research and innovation programme under the grant agreement ERC-2016-724173 the Marie Sklodowska-Curie grant agreement No H2020-MSCA-ITN-2014-642095. DT was funded by “Severo Ochoa visiting scientific programme” for a 6 months stay at the Center for Genomic Regulation - Barcelona - to start the project, supported as well by INRA SPE. Dr Roderic Guigo (CRG, Barcelona) is warmly acknowledged for his welcome in his lab.

Drs. Akiko Sugio, Julie Jaquiéry, Gaël Le Trionnaire and Jean-Christophe Simon (INRA, UMR 1349 Igepp, Rennes, France) are acknowledged for access to unpublished data of RNA-Seq. *Daktulosphaira vitifoliae* data were provided by the Phylloxera Genome Project (https://bipaa.genouest.org/is/aphidbase/): funding for *D. vitifoliae* clone Pcf genomic sequencing was provided by INRA (AIP Bioressources) and BGI Biotech in the frame of i5k initiative. Parts of the transcriptomic resources were obtained within the 1KITE projects (Bernhard Misof, Bonn, Germany).

The *Acyrthosiphon pisum* phylome can be accessed at PhylomeDB 4.0 under phylome number 441. RNA-Seq data are accessible at NCBI, see **Table S2**.

## Author contributions

RF designed research, analysed data, wrote the manuscript and prepared figures and tables. MMH designed research, analysed data, wrote the manuscript and prepared figures and tables. FL and SR performed bioinformatic analyses on RNA-Seq methods. GR produced FAIRE-Seq data and participated in the analysis and discussion of that section. VW was involved in the early steps of RNA-Seq analyses. CP designed research, analysed data, wrote the manuscript and prepared figures and tables. TG, DT designed and supervised research, coordinated the production of data and supervised the writing of the manuscript.

## References

Ackermann AM, Wang Z, Schug J, Naji A, Kaestner KH. 2016. Integration of ATAC-seq and RNA-seq identifies human alpha cell and beta cell signature genes. Mol Metab 5:233–244.

Al-Shahrour F, Diaz-Uriarte R, Dopazo J. 2004. FatiGO: a web tool for finding significant associations of Gene Ontology terms with groups of genes. Bioinformatics 20:578–580.

Athey J, Alexaki A, Osipova E, Rostovtsev A, Santana-Quintero LV, Katneni U, Simonyan V, Kimchi-Sarfaty C. 2017. A new and updated resource for codon usage tables. BMC Bioinformatics 18:391.

Capella-Gutierrez S, Silla-Martinez JM, Gabaldon T. 2009. trimAl: a tool for automated alignment trimming in large-scale phylogenetic analyses. Bioinformatics 25:1972–1973.

Chang AY-F, Liao B-Y. 2012. DNA methylation rebalances gene dosage after mammalian gene duplications. Mol. Biol. Evol. 29:133–144.

Churchman LS, Weissman JS. 2012. Native elongating transcript sequencing (NET-seq). Curr. Protoc. Mol. Biol. Chapter 4:Unit 4.14.1–17.

Conant GC. 2003. Asymmetric Sequence Divergence of Duplicate Genes. Genome Res. 13:2052–2058.

Dobin A, Davis CA, Schlesinger F, Drenkow J, Zaleski C, Jha S, Batut P, Chaisson M, Gingeras TR. 2013. STAR: ultrafast universal RNA-seq aligner. Bioinformatics 29:15–21.

Duncan RP, Feng H, Nguyen DM, Wilson ACC. 2016. Gene Family Expansions in Aphids Maintained by Endosymbiotic and Nonsymbiotic Traits. Genome Biol. Evol. 8:753–764.

Edgar RC. 2004. MUSCLE: multiple sequence alignment with improved accuracy and speed. Proceedings. 2004 IEEE Computational Systems Bioinformatics Conference, 2004. CSB 2004.

Etherington GJ, Ramirez-Gonzalez RH, MacLean D. 2015. bio-samtools 2: a package for analysis and visualization of sequence and alignment data with SAMtools in Ruby: Fig. 1. Bioinformatics 31:2565–2567.

Guindon S, Dufayard J-F, Lefort V, Anisimova M, Hordijk W, Gascuel O. 2010. New algorithms and methods to estimate maximum-likelihood phylogenies: assessing the performance of PhyML 3.0. Syst. Biol. 59:307–321.

Han MV, Demuth JP, McGrath CL, Casola C, Hahn MW. 2009. Adaptive evolution of young gene duplicates in mammals. Genome Res. 19:859–867.

Hansen AK, Moran NA. 2011. Aphid genome expression reveals host-symbiont cooperation in the production of amino acids. Proc. Natl. Acad. Sci. U. S. A. 108:2849–2854.

Huerta-Cepas J, Capella-Gutierrez S, Pryszcz LP, Denisov I, Kormes D, Marcet-Houben M, Gabaldón T. 2011. PhylomeDB v3.0: an expanding repository of genome-wide collections of trees, alignments and phylogeny-based orthology and paralogy predictions. Nucleic Acids Res. 39:D556–D560.

Huerta-Cepas J, Capella-Gutiérrez S, Pryszcz LP, Marcet-Houben M, Gabaldón T. 2014. PhylomeDB v4: zooming into the plurality of evolutionary histories of a genome. Nucleic Acids Res. 42:D897–D902.

Huerta-Cepas J, Dopazo J, Gabaldón T. 2010. ETE: a python Environment for Tree Exploration. BMC Bioinformatics 11.

Huerta-Cepas J, Dopazo J, Huynen MA, Gabaldón T. 2011. Evidence for short-time divergence and long-time conservation of tissue-specific expression after gene duplication. Brief. Bioinform. 12:442–448.

Huerta-Cepas J, Forslund K, Coelho LP, Szklarczyk D, Jensen LJ, von Mering C, Bork P. 2017. Fast Genome-Wide Functional Annotation through Orthology Assignment by eggNOG-Mapper. Molecular Biology and Evolution 34:2115–2122.

Huerta-Cepas J, Marcet-Houben M, Pignatelli M, Moya A, Gabaldón T. 2010. The pea aphid phylome: a complete catalogue of evolutionary histories and arthropod orthology and paralogy relationships for Acyrthosiphon pisum genes. Insect Mol. Biol. 19 Suppl 2:13–21.

Huerta-Cepas J, Szklarczyk D, Heller D, Hernández-Plaza A, Forslund SK, Cook H, Mende DR, Letunic I, Rattei T, Jensen LJ, et al. 2019. eggNOG 5.0: a hierarchical, functionally and phylogenetically annotated orthology resource based on 5090 organisms and 2502 viruses. Nucleic Acids Research 47:D309–D314.

Innan H, Kondrashov F. 2010. The evolution of gene duplications: classifying and distinguishing between models. Nat. Rev. Genet. 11:97–108.

Kanayama HO, Tamura T, Ugai S, Kagawa S, Tanahashi N, Yoshimura T, Tanaka K, Ichihara A. 1992. Demonstration that a human 26S proteolytic complex consists of a proteasome and multiple associated protein components and hydrolyzes ATP and ubiquitin-ligated proteins by closely linked mechanisms. Eur. J. Biochem. 206:567–578.

Katoh K. 2005. MAFFT version 5: improvement in accuracy of multiple sequence alignment. Nucleic Acids Research 33:511–518.

Keller TE, Yi SV. 2014. DNA methylation and evolution of duplicate genes. Proceedings of the National Academy of Sciences 111:5932–5937.

Kofler R, Orozco-terWengel P, De Maio N, Pandey RV, Nolte V, Futschik A, Kosiol C, Schlötterer C. 2011. PoPoolation: a toolbox for population genetic analysis of next generation sequencing data from pooled individuals. PLoS One 6:e15925.

Kumar S, Stecher G, Suleski M, Hedges SB. 2017. TimeTree: A Resource for Timelines, Timetrees, and Divergence Times. Mol. Biol. Evol. 34:1812–1819.

Langmead B, Salzberg SL. 2012. Fast gapped-read alignment with Bowtie 2. Nat. Methods 9:357–359.

Langmead B, Trapnell C, Pop M, Salzberg SL. 2009. Ultrafast and memory-efficient alignment of short DNA sequences to the human genome. Genome Biol 10: R25.

Lassmann T, Sonnhammer ELL. 2005. Kalign--an accurate and fast multiple sequence alignment algorithm. BMC Bioinformatics 6:298.

Le SQ, Lartillot N, Gascuel O. 2008. Phylogenetic mixture models for proteins. Philos. Trans. R. Soc. Lond. B Biol. Sci. 363:3965–3976.

Liao Y, Smyth GK, Shi W. 2014. featureCounts: an efficient general purpose program for assigning sequence reads to genomic features. Bioinformatics 30:923–930.

Li H, Handsaker B, Wysoker A, Fennell T, Ruan J, Homer N, Marth G, Abecasis G, Durbin R, 1000 Genome Project Data Processing Subgroup. 2009. The Sequence Alignment/Map format and SAMtools. Bioinformatics 25:2078–2079.

Li Q, Brown JB, Huang H, Bickel PJ. 2011. Measuring reproducibility of high-throughput experiments. Ann. Appl. Stat. 5:1752–1779.

Li Y, Park H, Smith TE, Moran NA. 2019. Gene Family Evolution in the Pea Aphid Based on Chromosome-Level Genome Assembly. Mol. Biol. Evol. 36:2143–2156.

Lynch M, Conery JS. 2000. The evolutionary fate and consequences of duplicate genes. Science 290:1151–1155.

Mansai SP, Kado T, Innan H. 2011. The Rate and Tract Length of Gene Conversion between Duplicated Genes. Genes 2:313–331.

Marlétaz F, Firbas PN, Maeso I, Tena JJ, Bogdanovic O, Perry M, Wyatt CDR, de la Calle-Mustienes E, Bertrand S, Burguera D, et al. 2018. Amphioxus functional genomics and the origins of vertebrate gene regulation. Nature 564:64–70.

Mathers TC, Chen Y, Kaithakottil G, Legeai F, Mugford ST, Baa-Puyoulet P, Bretaudeau A, Clavijo B, Colella S, Collin O, et al. 2017. Rapid transcriptional plasticity of duplicated gene clusters enables a clonally reproducing aphid to colonise diverse plant species. Genome Biol. 18:27.

McCarthy DJ, Chen Y, Smyth GK. 2012. Differential expression analysis of multifactor RNA-Seq experiments with respect to biological variation. Nucleic Acids Res. 40:4288–4297.

Mirarab S, Nguyen N, Guo S, Wang L-S, Kim J, Warnow T. 2015. PASTA: Ultra-Large Multiple Sequence Alignment for Nucleotide and Amino-Acid Sequences. J. Comput. Biol. 22:377–386.

Pegueroles C, Laurie S, Albà MM. 2013. Accelerated evolution after gene duplication: a time-dependent process affecting just one copy. Mol. Biol. Evol. 30:1830–1842.

Pich I Roselló O, Kondrashov FA. 2014. Long-term asymmetrical acceleration of protein evolution after gene duplication. Genome Biol. Evol. 6:1949–1955.

Puigbò P, Bravo IG, Garcia-Vallve S. 2008. CAIcal: a combined set of tools to assess codon usage adaptation. Biol. Direct 3:38.

Rajaiya J, Nixon JC, Ayers N, Desgranges ZP, Roy AL, Webb CF. 2006. Induction of immunoglobulin heavy-chain transcription through the transcription factor Bright requires TFII-I. Mol. Cell. Biol. 26:4758–4768.

Ramírez F, Ryan DP, Grüning B, Bhardwaj V, Kilpert F, Richter AS, Heyne S, Dündar F, Manke T. 2016. deepTools2: a next generation web server for deep-sequencing data analysis. Nucleic Acids Res. 44:W160–W165.

Reddy VS, Shlykov MA, Castillo R, Sun EI, Saier MH. 2012. The major facilitator superfamily (MFS) revisited. FEBS Journal 279:2022–2035.

Rice P, Craigie R, Davies DR. 1996. Retroviral integrases and their cousins. Curr. Opin. Struct. Biol. 6:76–83.

Richard G. 2017. Régulations chromatiniennes et transcriptionnelles impliquées dans le cycle de vie du puceron du pois. Available from: http://www.theses.fr/2017NSARB130

Richard G, Legeai F, Prunier-Leterme N, Bretaudeau A, Tagu D, Jaquiéry J, Le Trionnaire G. 2017. Dosage compensation and sex-specific epigenetic landscape of the X chromosome in the pea aphid. Epigenetics Chromatin 10:30.

Robinson MD, McCarthy DJ, Smyth GK. 2010. edgeR: a Bioconductor package for differential expression analysis of digital gene expression data. Bioinformatics 26:139–140.

Sandve SR, Rohlfs RV, Hvidsten TR. 2018. Subfunctionalization versus neofunctionalization after whole-genome duplication. Nat. Genet. 50:908–909.

Sato K, Nishida KM, Shibuya A, Siomi MC, Siomi H. 2011. Maelstrom coordinates microtubule organization during Drosophila oogenesis through interaction with components of the MTOC. Genes Dev. 25:2361–2373.

Sawyer S. 1989. Statistical tests for detecting gene conversion. Mol. Biol. Evol. 6:526–538.

Scannell DR, Wolfe KH. 2008. A burst of protein sequence evolution and a prolonged period of asymmetric evolution follow gene duplication in yeast. Genome Res. 18:137–147.

Simon J-C, Pfrender ME, Tollrian R, Tagu D, Colbourne JK. 2011. Genomics of environmentally induced phenotypes in 2 extremely plastic arthropods. J. Hered. 102:512–525.

Simon JM, Giresi PG, Davis IJ, Lieb JD. 2012. Using formaldehyde-assisted isolation of regulatory elements (FAIRE) to isolate active regulatory DNA. Nature Protocols 7:256–267.

Supek F, Bošnjak M, Škunca N, Šmuc T. 2011. REVIGO Summarizes and Visualizes Long Lists of Gene Ontology Terms. PLoS One 6:e21800.

Takahashi H, Kato S, Murata M, Carninci P. 2012. CAGE (cap analysis of gene expression): a protocol for the detection of promoter and transcriptional networks. Methods Mol. Biol. 786:181–200.

The International Aphid Genomics Consortium. 2010. Genome Sequence of the Pea Aphid Acyrthosiphon pisum. PLoS Biol. 8:e1000313.

Vellichirammal NN, Madayiputhiya N, Brisson JA. 2016. The genomewide transcriptional response underlying the pea aphid wing polyphenism. Mol. Ecol. 25:4146–4160.

Venkat A, Hahn MW, Thornton JW. 2018. Multinucleotide mutations cause false inferences of lineage-specific positive selection. Nat Ecol Evol 2:1280–1288.

Wallace IM, O’Sullivan O, Higgins DG, Notredame C. 2006. M-Coffee: combining multiple sequence alignment methods with T-Coffee. Nucleic Acids Res. 34:1692–1699.

Wehe A, Bansal MS, Burleigh JG, Eulenstein O. 2008. DupTree: a program for large-scale phylogenetic analyses using gene tree parsimony. Bioinformatics 24:1540–1541.

Werner MS, Sieriebriennikov B, Prabh N, Loschko T, Lanz C, Sommer RJ. 2018. Young genes have distinct gene structure, epigenetic profiles, and transcriptional regulation. Genome Res. 28:1675–1687.

Yang Z. 2007. PAML 4: phylogenetic analysis by maximum likelihood. Mol. Biol. Evol. 24:1586–1591.

Zhang J. 2005. Evaluation of an Improved Branch-Site Likelihood Method for Detecting Positive Selection at the Molecular Level. Molecular Biology and Evolution 22:2472–2479.

Zhang Y, Liu T, Meyer CA, Eeckhoute J, Johnson DS, Bernstein BE, Nusbaum C, Myers RM, Brown M, Li W, et al. 2008. Model-based analysis of ChIP-Seq (MACS). Genome Biol. 9:R137.

