## Supplementary material for "Selection following gene duplication shapes recent genome evolution in the pea aphid *Acyrthosiphon pisum*": Suppkemental Figures (from another version of this preprint: version 1, 2019-05-20)

**Supplemental Figures.**

**Figure S1.** Median similarities for each protein sequence alignment for the selected duplications after filtering duplications with dS >2 and dS <0.01. P-values were estimated using the `wilcox.test` function from R. A: *A. pisum* genes from the 572 selected duplications. B: *A. pisum* genes were divided into two categories: those having signals of positive selection (PS=1, depicted in yellow) and those with no signals of positive selection (PS=0, depicted in green).

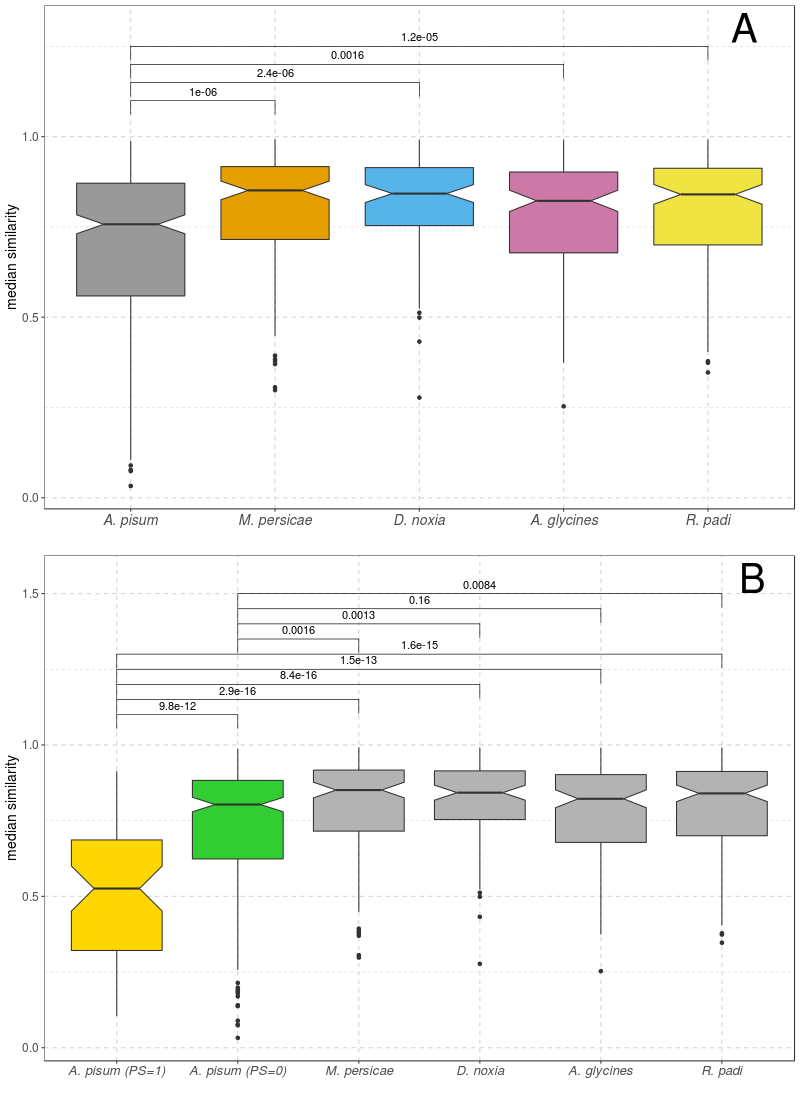

**Figure S2**. Treemap of GO enrichment summary of the 1,144 duplicated genes selected for further analyses, as described in the main text. Square size is proportional to GO term presence and hierarchical relationships between GO levels. This dense map indicates the large number of functions covered by the selected duplicated genes. See full annotation results in Table S8.

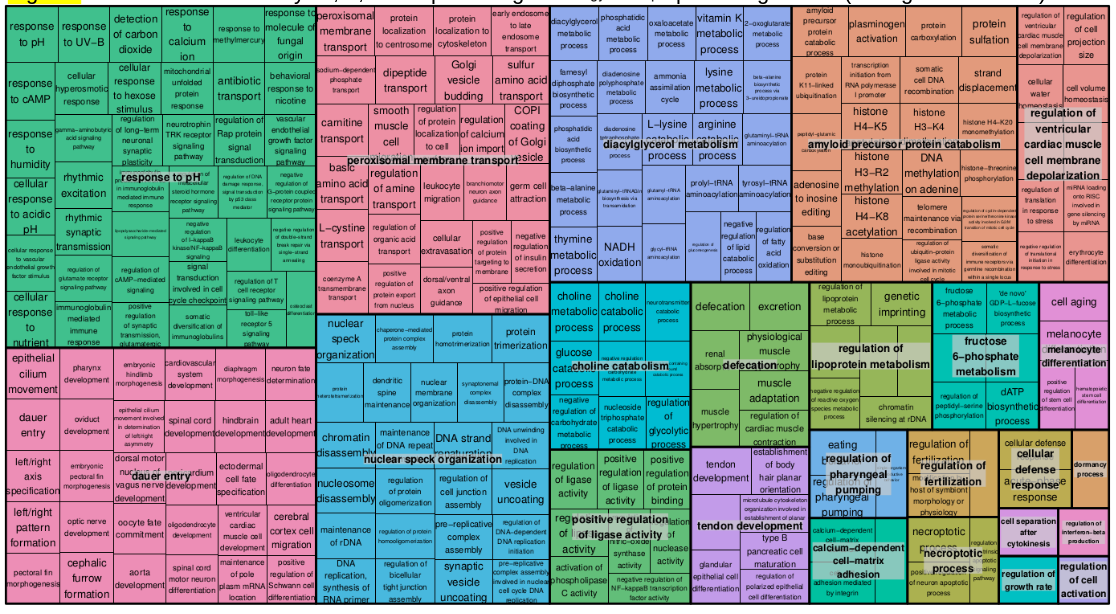

**Figure S3.** Treemap of GO terms illustrating the multiple functions of all genes under selection. Since this is not an enrichment analysis, square size is proportional to GO term presence and hierarchical relationships between GO levels. See full annotation results in Table S9.

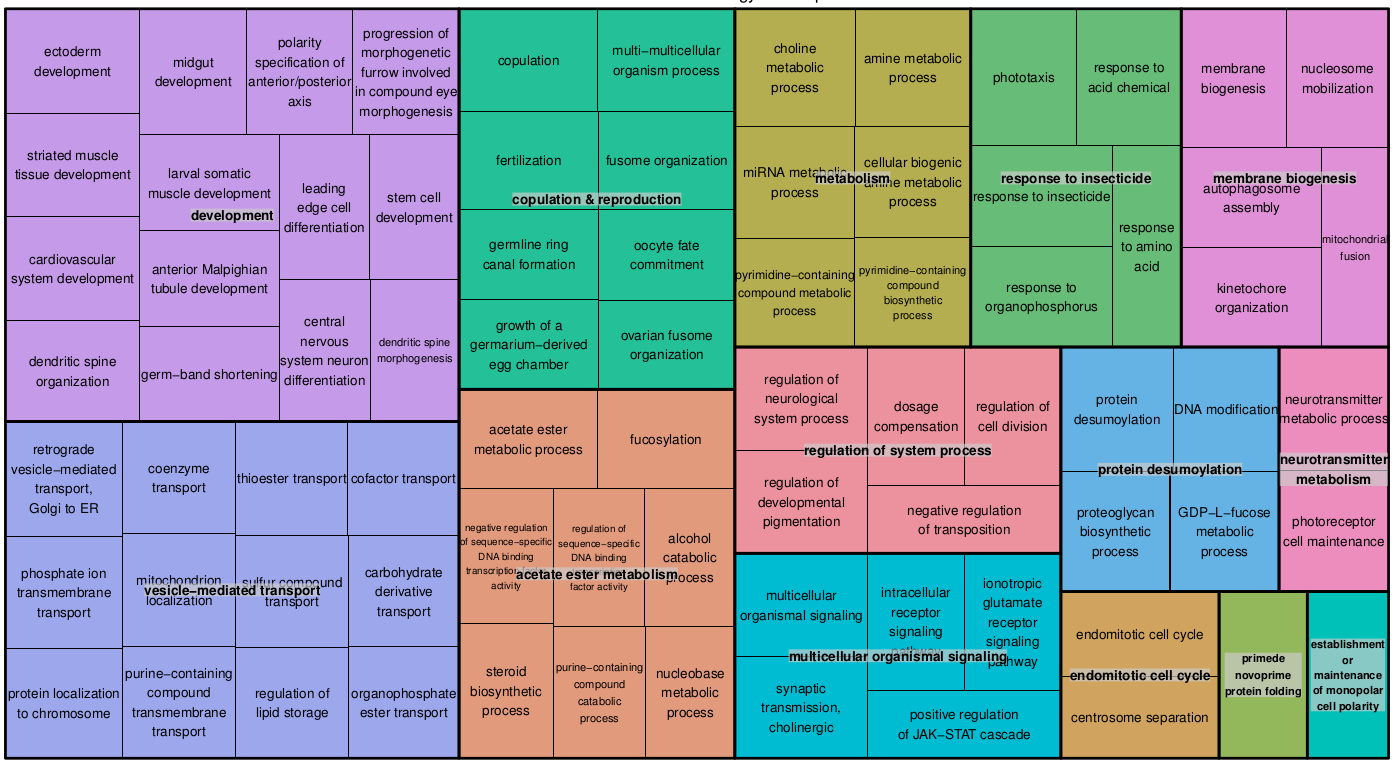

**Figure S4:** Biologically-relevant GO terms and BLAST annotations of some of the positively-selected genes.

a) ***Resistance to insecticides***

**GO:0017085 -> resistance to insecticide**

Gene ID: Phy00BWVM0_ACYPI-> BLAST annotation: aromatic-L-amino-acid decarboxylase

Gene ID: Phy00423PN_ACYPI -> BLAST annotation: ecdysone 20-monooxygenase isoform X1

***b) Immunology***

**GO:0002920 regulation of humoral immune response**

**GO:0002805 regulation of antimicrobial peptide biosynthetic process**

**GO:0002784 regulation of antimicrobial peptide production**

**GO:0002759 regulation of antimicrobial humoral response**

Gene ID: Phy00BX1FX_ACYPI -> BLAST annotation: serine protease inhibitor 4, serpin-4-like isoform X1

Gene ID: Phy00BWVQ3_ACYPI -> BLAST annotation: zinc finger protein ush-like

***c) Reproduction***

**GO:2000035 regulation of stem cell division**

Gene ID: Phy004221P_ACYPI -> BLAST annotation: GDP-mannose 4,6 dehydratase-like isoform X1

**GO:0048132 female germ-line stem cell asymmetric division**

Gene ID: Phy00BWWQD_ACYPI -> BLAST annotation: interferon-inducible double-stranded RNA-dependent protein kinase activator A-like

**GO:0030718 germ-line stem cell population maintenance**

Gene ID: Phy00BWWQD_ACYPI -> BLAST annotation: interferon-inducible double-stranded RNA-dependent protein kinase activator A-like

Gene ID: Phy00BX2G6_ACYPI -> BLAST annotation: ubiquitin carboxyl-terminal hydrolase 36-like

Gene ID: Phy000YNOH_ACYPI -> BLAST annotation: protein maelstrom homolog

Gene ID: Phy00429XW_ACYPI -> BLAST annotation: protein maelstrom 2

**GO:0001700 embryonic development via the syncytial blastoderm**

Gene ID: Phy00423PN_ACYPI -> BLAST annotation: ecdysone 20-monooxygenase isoform X1 Gene ID: Phy00BWVQ3_ACYPI -> BLAST annotation: zinc finger protein ush-like

Gene ID: Phy00421YB_ACYPI -> BLAST annotation: zinc finger protein 865-like

**GO:0030716 oocyte fate determination**

Gene ID: Phy00BWZ4J_ACYPI -> BLAST annotation: helicase POLQ-like

Gene ID: Phy00BWW2V_ACYPI-> BLAST annotation: helicase POLQ-like

**GO:0048600 oocyte fate commitment**

Gene ID: Phy000YNOH_ACYPI -> BLAST annotation: protein maelstrom homolog

Gene ID: Phy00429XW_ACYPI -> BLAST annotation: protein maelstrom 2

**GO:0007350 blastoderm segmentation**

**GO:0007354 zygotic determination of anterior/posterior axis, embryo**

**GO:0008358 maternal determination of anterior/posterior axis, embryo**

Gene ID: Phy00BX3PU_ACYPI -> BLAST annotation: bifunctional heparan sulfate N-deacetylase/N-sulfotransferase

Gene ID: Phy00BWVQ3_ACYPI-> BLAST annotation: zinc finger protein ush-like

Gene ID: Phy0041ZGT_ACYPI -> BLAST annotation: E3 ubiquitin-protein ligase TRIM33-likeGene ID: Phy000YNOH_ACYPI -> BLAST annotation: protein maelstrom homolog

Gene ID: Phy00429XW_ACYPII -> BLAST annotation: protein maelstrom 2

**GO:0007300 ovarian nurse cell to oocyte transport**

Gene ID: Phy00BWYDK_ACYPI -> BLAST annotation: kelch-like protein 2

Gene ID: Phy00420VZ_ACYPI -> BLAST annotation: kelch-like protein 2

**GO:0016321 female meiosis chromosome segregation**

Gene ID: Phy00BWZ4J_ACYPI -> BLAST annotation: helicase POLQ-like

Gene ID: Phy00BWW2V_ACYPI-> BLAST annotation: helicase POLQ-like

**GO:00030717 karyosome formation**

**GO:0016325 oocyte microtubule cytoskeleton organization**

**GO:0007293 germarium-derived egg chamber formation**

**GO:0007308 oocyte construction**

Gene ID: Phy00BWZ4J_ACYPI -> BLAST annotation: helicase POLQ-like

Gene ID: Phy00BWW2V_ACYPI-> BLAST annotation: helicase POLQ-like

Gene ID: Phy00BWYDK_ACYPI -> BLAST annotation: kelch-like protein 2

Gene ID: Phy00420VZ_ACYPI -> BLAST annotation: kelch-like protein 2

**GO:0030725 germline ring canal formation**

Gene ID: Phy00BWYDK_ACYPI-> BLAST annotation: kelch-like protein 2

Gene ID: Phy00420VZ_ACYPI -> BLAST annotation: kelch-like protein 2

***d) Morphogenesis & Development***

**GO:0048069 eye pigmentation**

Gene ID: Phy00BWVCW_ACYPI -> BLAST annotation: peptidyl-prolyl cis-trans isomerase 5-like

**GO:0007398 ectoderm development**

Gene ID: Phy00BWVQ3_ACYPI-> BLAST annotation: zinc finger protein ush-like

**GO:0007458 progression of morphogenetic furrow involved in compound eye morphogenesis**

Gene ID: Phy00421YB_ACYPI-> BLAST annotation: zinc finger protein 865-like

**GO:0007525 somatic muscle development**

Gene ID: Phy00BX0AY_ACYPI -> BLAST annotation: cGMP-dependent protein kinase 1

Gene ID: Phy00BWVDL_ACYPI -> BLAST annotation: cGMP-dependent protein kinase, isozyme 1

**GO:0008594 photoreceptor cell morphogenesis**

Gene ID: Phy00BWVDL_ACYPI -> BLAST annotation: cGMP-dependent protein kinase, isozyme 1

**GO:0007494 midgut development**

Gene ID: Phy00423PN_ACYPI -> BLAST annotation: ecdysone 20-monooxygenase isoform X1

**GO:0065001 specification of axis polarity**

Gene ID: Phy00BWZ4J_ACYPI -> BLAST annotation: helicase POLQ-like

Gene ID: Phy00BWW2V_ACYPI -> BLAST annotation: helicase POLQ-like

**GO:0072002 Malpighian tubule development**

Gene ID: Phy00BX35W_ACYPI -> BLAST annotation: protein TANC1

Gene ID: Phy00BX2RE_ACYPI -> BLAST annotation: protein TANC2 isoform X1

Gene ID: Phy0042A0T_ACYPI -> BLAST annotation: collagen alpha-1(IV) chain-like

***d) Dosage compensation***

**GO:0009047->** dosage compensation by hypoactivation of X chromosome

**GO:0006344** -> maintenance of chromatin silencing

Gene ID: Phy00BWXWJ_ACYPI -> BLAST annotation: male-specific lethal 3 homolog

**GO:0060966 regulation of gene silencing by RNA**

**GO:0060964 regulation of gene silencing by miRNA**

**GO:0060147 regulation of postranscriptional gene silencing**

Gene ID: Phy00BWWQD_ACYPI -> BLAST annotation: interferon-inducible double-stranded RNA-dependent protein kinase activator A-like

Gene ID: Phy000YNOH_ACYPI -> BLAST annotation: protein maelstrom homolog

Gene ID: Phy00429XW_ACYPII -> BLAST annotation: protein maelstrom 2

***e) Ecdysis and metamorphosis***

**GO:0045456 ecdysteroid biosynthetic process**

Gene ID: Phy00423PN_ACYPI -> BLAST annotation: ecdysone 20-monooxygenase isoform X1

**GO:0040003 chitin-based cuticle development**

**GO:0048067 cuticle pigmentation**

**GO:0007564 regulation of chitin-based cuticle tanning**

Gene ID: Phy00BWY9R_ACYPI -> BLAST annotation: low-density lipoprotein receptor-related protein 2

**GO:0042335 cuticle development**

Gene ID: Phy00BWY9R_ACYPI

Gene ID: Phy004203L_ACYPI -> BLAST annotation: protein yellow-like

**GO:0022416 chaeta development**

Gene ID: Phy00BX50P_ACYPI -> BLAST annotation: baculoviral IAP repeat-containing protein 3 Gene ID: Phy00BWVQ3_ACYPI-> BLAST annotation: zinc finger protein ush-like

**GO:0035075 response to ecdysone**

**GO:0036315 cellular response to sterol**

Gene ID: Phy0041ZOV_ACYPI -> BLAST annotation: inorganic pyrophosphatase

**Figure S5.** Scatterplots comparing the three different statistics to evaluate differences of gene expression across tissues: Hamming distance, tissue expression complementarity (TEC) distance and tissue divergence (dT). Pearson correlation values are provided for each comparison (top right corner of each panel).

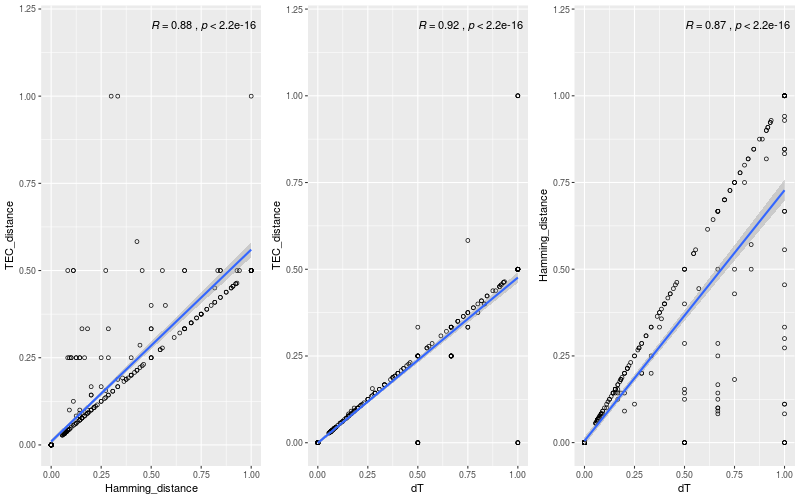

**Figure S6.** Tissue expression differences for duplications that have different expression patterns in at least one of the studied tissues. This duplications have been classified in cases with no positive selection (PS=0), positive selection in one copy (PS=1) and positive selection in the two copies (PS=2, merely one case). We used three different statistics to evaluate differences of gene expression across tissues: Hamming distance, tissue expression complementarity (TEC) distance and tissue divergence (dT).

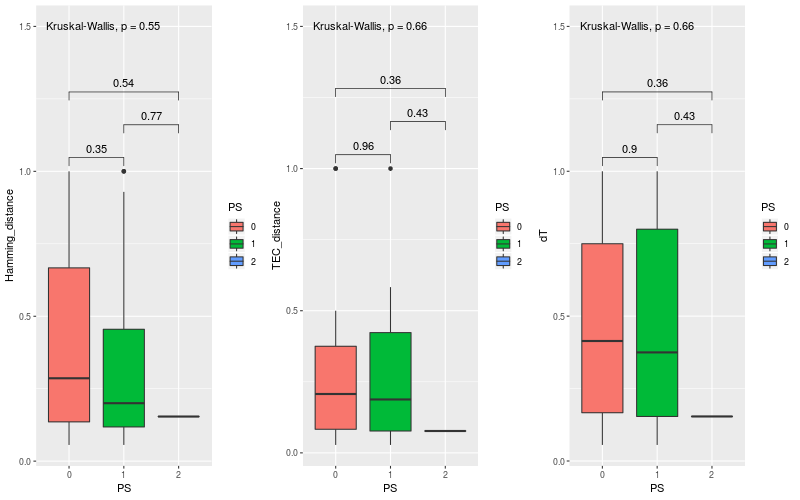
